# Pollen tube-triggered accumulation of NORTIA at the filiform apparatus facilitates fertilization in *Arabidopsis thaliana*

**DOI:** 10.1101/621599

**Authors:** Jing Yuan, Yan Ju, Daniel S. Jones, Weiwei Zhang, Noel Lucca, Christopher J. Staiger, Sharon A. Kessler

## Abstract

During gamete delivery in *Arabidopsis thaliana*, intercellular communication between the attracted pollen tube and the receptive synergid cell leads to subcellular events in both cells culminating in the rupture of the tip-growing pollen tube and release of the sperm cells to achieve double fertilization. Live imaging of pollen tube reception revealed dynamic subcellular changes that occur in the female synergid cells. Pollen tube arrival triggers the trafficking of NORTIA (NTA) MLO protein from Golgi-associated compartments and the accumulation of endosomes at or near the synergid filiform apparatus, a membrane-rich region that acts as the site of communication between the pollen tube and synergids. Domain swaps and site-directed mutagenesis reveal that NTA’s C-terminal cytoplasmic tail with its calmodulin-binding domain influences the subcellular localization and function of NTA in pollen tube reception and that accumulation of NTA at the filiform apparatus is necessary and sufficient for MLO function in pollen tube reception.

## Introduction

Intercellular communication is central to the proper development and maintenance of all multicellular organisms. During this communication, signals from one cell are perceived by receptors in another cell and translated into various subcellular responses. These responses include signal transduction cascades leading to transcription of other genes, calcium signaling, and trafficking of proteins to different organelles or regions of the cell. A well-studied example of signal-induced protein trafficking in plant development is the redistribution of the PIN polar auxin transporters to different sides of the cell during important developmental events such as embryo patterning, leaf initiation and lateral root initiation [1–3]. In plants, most intercellular communication occurs between cells that are genetically identical and connected by adjoining cell walls. One exception is pollination, in which pollen (the male gametophyte) is released from an anther, transported to a receptive stigma, and produces a tip-growing pollen tube that grows through the female tissues of the pistil and delivers the two sperm cells to the female gametophyte (also known as the embryo sac, Figure 1A). The pollen tube’s journey through the pistil requires cell-to-cell interactions with the female that allows water and nutrient uptake and enables the detection of cues important for guidance toward the female gametes [4].

**Figure 1.**
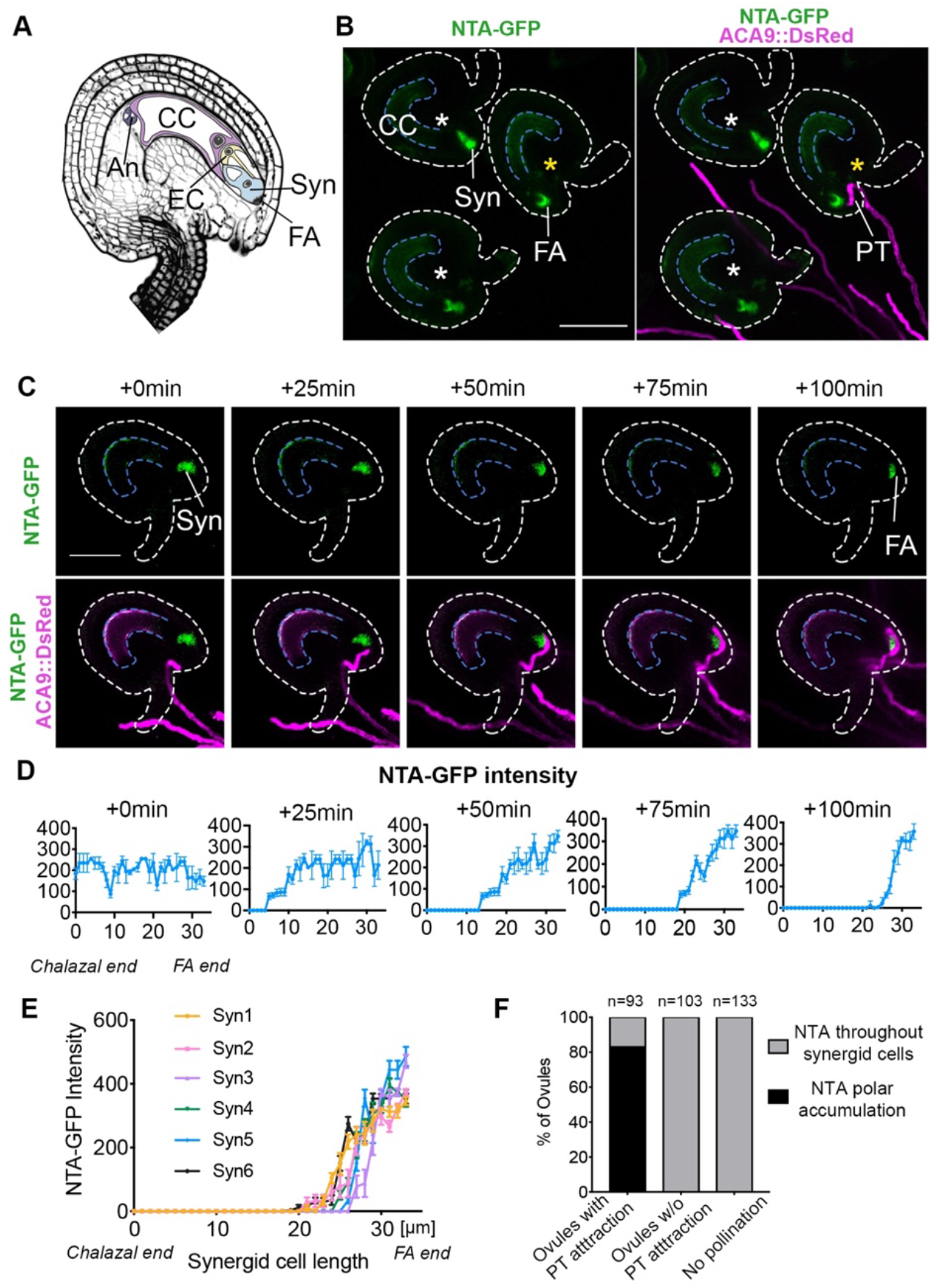
NTA-GFP accumulates at the filiform apparatus as the pollen tube approaches. (A) Diagram of a mature *Arabidopsis thaliana* ovule and embryo sac, modified from Jones et al, 2018. CC, Central Cell; Syn, Synergid Cells; EC, Egg Cell; An, Antipodal cells; FA, Filiform Apparatus (B, C) Live imaging of pollen tube (PT) reception using NTA-GFP labeled synergids (green signal) and *ACA9::DsRed* pollen tubes (magenta signal). (B) Polar NTA-GFP accumulation at the filiform apparatus (FA) occurred in ovules that attracted a pollen tube (ovules with yellow stars), while polar NTA-GFP accumulation did not occur in ovules without pollen tube attraction (ovules with white stars). (C) Time-lapse imaging of NTA-GFP accumulation during pollen tube reception. (D) Quantification of NTA-GFP signal along the length of the synergid from chalazal end (0μm) to FA end (33μm) at 0min, 25min, 50min, 75min, and 100min timepoints, respectively. (E) 6 more examples of the quantification of NTA-GFP signal along the length of synergids after pollen tube arrival. (F) Quantification of the percentage of ovules with different NTA accumulation patterns under the same imaging conditions. Bars=50 μm (B and C).

In the model plant *Arabidopsis thaliana*, complex signaling events ranging from pollen landing on the stigma to fusion of gametes occur over several hours. Most of our knowledge about the signaling pathways involved along the pollen tube’s journey through the female is limited to the final stages of pollination and involve a highly specialized pair of female gametophyte cells known as synergids. During female gametophyte development, meiosis followed by three rounds of mitosis produce the egg cell and central cell along with 2 synergid cells flanking the egg cell and 3 antipodal cells on the chalazal end of the embryo sac [5] (Figure 1A). The synergid cells are accessory cells that control the behavior of the pollen tube during the final stages of pollination. Before pollen tube arrival, they secrete cysteine-rich LURE peptides that act as short-range pollen tube attractants that are recognized by receptor-like kinases in the tip of the pollen tube to regulate the direction of pollen tube growth and guide the pollen tube to the micropyle of the ovule [6–8]. After pollen tube arrival, the synergids communicate with the pollen tube to induce changes that result in pollen tube rupture and delivery of the sperm cells [4, 9]. Thus, synergids are critical for ensuring that double fertilization can occur to produce seeds.

In Arabidopsis, live imaging has been used to examine the behavior of both the pollen tube and the synergids during the process of pollen tube reception. A pollen tube follows the gradient of LURE attractants, enters the micropyle of the ovule, and pauses its growth for 30 min to 1 h just outside the receptive synergid [10–12]. During this pause in pollen tube growth, communication occurs between the pollen tube and the synergids leading to subcellular changes and ultimately the death of both the pollen tube and the receptive synergid. Cytoplasmic calcium oscillations occur in both the tip of the pollen tube and in the 2 synergid cells during this communication phase. Cytoplasmic calcium levels continue to increase in both cell types until the pollen tube starts to grow again and bursts to release the sperm cells, a catastrophic event for both the pollen tube and the receptive synergid, which also degenerates [10–12]. Mutations in genes that regulate communication between the synergids and pollen tube during pollen tube reception result in a pollen tube overgrowth phenotype in which the pollen tubes are attracted normally to the ovules, but do not get the signal to burst and release the sperm cells. Presumably, synergid-induced changes in the cell wall of the pollen tube tip do not occur in these mutants, therefore the pollen tube continues to grow and coil inside the embryo sac. Synergid-expressed genes that participate in pollen tube reception include the FERONIA (FER) receptor-like kinase, the GPI-anchored protein LORELEI (LRE), and the Mildew Resistance Locus-O (MLO) protein NORTIA (NTA, also known as AtMLO7) [12–17]. Mutations in all of these genes lead to the pollen tube overgrowth phenotype due to disruption of the pollen tube-synergid communication pathway.

FER and LRE are necessary for the calcium oscillations that occur in synergids in response to pollen tube arrival [12]. In contrast, *nta-1* mutants have [Ca^2+^]_cyto_ oscillations at lower amplitudes, indicating that NTA may participate in modulating Ca^2+^ fluxes in the synergids during communication with the pollen tube and likely act downstream of FER and LRE [12]. Like all members of the MLO gene family, NTA has seven membrane-spanning domains and a predicted calmodulin-binding domain (CaMBD) in its C-terminal intracellular tail [18, 19]. Calmodulin (CaM) is a small protein that binds Ca^2+^ and is involved in signal transduction for many cellular processes [20]. We previously showed that the C-terminal domain of NTA could confer pollen tube reception function to the related MLO8 protein [21], but the significance of the CaMBD in pollen tube reception remains an open question.

The subcellular localization of important pollen tube reception proteins is not always predictive of their function in communicating with the pollen tube. As expected for early response proteins, both FER and LRE are expressed in synergid cells where they localize in or near a specialized region called the filiform apparatus, a membrane rich area located at the micropyle end of the synergids [14–17, 22–24]. The filiform apparatus is thought to be important for the secretion of attractant peptides and is the first site of interaction between the pollen tube and synergid cell prior to pollen tube reception [25–27]. In contrast, before pollen tube arrival, NTA is localized to a Golgi-associated compartment within the synergid cell and absent from the filiform apparatus [21]. At the end of pollen tube reception, NTA protein is only detected at the filiform apparatus, indicating that this protein changes its subcellular localization during pollen tube reception [13]. This suggests that pollen tube-triggered regulation of the synergid secretory system may be a crucial subcellular response to pollen tube arrival and that NTA function may be related to its subcellular distribution; however, the precise timing and significance of NTA’s redistribution remain unclear.

Here, we use a live-imaging system to further characterize synergid cellular dynamics during pollen tube reception and to determine the timing and significance of the polar redistribution of NTA to the filiform apparatus. To investigate the link between Ca^2+^ and MLO function in pollen tube reception, we assayed the influence of the CaMBD on NTA’s function and subcellular distribution through C-terminal truncations and a point mutation disrupting the CaMBD. We show that the polar redistribution of NTA is triggered by the approach of a pollen tube and is facilitated by the CaMBD. While most subcellular compartments remain distributed throughout the synergid cells during pollen tube reception, recycling endosomes respond to pollen tube arrival by accumulating towards the filiform apparatus. Manipulation of NTA subcellular localization with domain swaps and a Golgi retention signal revealed that filiform apparatus localization is necessary and sufficient for MLO function in pollen tube reception.

## Results

### NTA dynamically accumulates at the filiform apparatus during pollen tube reception

Pollen tube reception requires synergid cells to recognize the approaching pollen tube and to send signals back to the pollen tube that result in release of the sperm cells at the correct time and place so that double fertilization can be completed. Based on static images, we previously reported that NTA-GFP fusion protein localizes to a Golgi-associated compartment in synergids prior to pollen tube attraction [21]. When imaged after pollen tube reception, NTA-GFP is concentrated at the micropylar end of the synergid (in or near the filiform apparatus) [13]. NTA-GFP does not accumulate at the filiform apparatus in *fer* ovules with pollen tube overgrowth, suggesting that FER-mediated signaling during pollen tube reception triggers NTA-GFP redistribution that in turn contributes to the interaction of the synergid with the pollen tube [13]. An alternative hypothesis is that pollen tube rupture triggers NTA-GFP redistribution and is a symptom of pollen tube reception rather than an important contributor to the signaling pathway. To distinguish between these two possibilities, we used a semi-*in vivo* pollination system combined with time-lapse spinning disk confocal microscopy to determine the timing of NTA-GFP redistribution during the pollen tube reception process. In the semi-*in vivo* system, pollen tubes grow out of a cut style and are attracted to ovules arranged on pollen germination media [28]. This system has previously been used to quantify and track pollen tube attraction to ovules and to image [Ca^2+^]_cyto_ reporters during pollen tube reception [10, 12, 29–32]. To follow subcellular changes in NTA-GFP protein localization before, during, and after pollen tube arrival, we used pollen from plants expressing the pollen-specific *AUTOINHIBITED Ca*^*2+−*^ *ATPase9*_*pro*_*::DsRed* (*ACA9_pro_::DsRed*) reporter and ovules expressing *NTA*_*pro*_*::NTA-GFP* in the semi-*in vivo* system. Approximately 4 h after pollination, pollen tubes emerged from the style onto the media and were attracted to ovules (Figure 1B). Fluorescence images in both channels were collected every 5 min from when a pollen tube approached an ovule until after the pollen tube ruptured inside the ovule. In ~83% (n=93) of the ovules that attracted a pollen tube and successfully burst to deliver the sperm cells, NTA-GFP accumulated at the filiform apparatus of the synergids (Figures 1C-E). The rest of the ovules (~17%) attracted pollen tubes that stopped growing in the micropyle and did not rupture. In these ovules, NTA-GFP did not accumulate <u>at the filiform</u> apparatus. To exclude the possibility that prolonged imaging causes stress in synergids which leads to filiform apparatus accumulation of NTA-GFP, we analyzed neighboring ovules that did not attract pollen tubes but were imaged together with ovules that attracted pollen tubes. NTA-GFP did not accumulate at the filiform apparatus in these ovules (n=103) (Figure 1F). Likewise, ovules that were incubated on pollen germination media without a pollinated pistil (n=133, Figures 1F and S1) and imaged over the same time frame did not accumulate NTA-GFP <u>at the filiform</u> apparatus. These data indicate that pollen tube arrival is necessary for NTA-GFP accumulation at the filiform apparatus rather than being retained in the Golgi, and that this accumulation is not an artifact of the semi-*in vivo* imaging system. (Figure 1F).

Our semi-*in vivo* system also allowed us to determine the timing of NTA-GFP accumulation at the filiform apparatus in relation to the position of the pollen tube as it approached the synergids. We defined the 0 min timepoint as the time where the pollen tube just reached the micropylar opening of the ovule (Figure 1C, ovules with yellow stars, Movies S1 and S2). In all cases, the shift of NTA-GFP signal from the Golgi to the filiform apparatus also started from this time point. During the following 30–50 min, pollen tubes grew through the micropyle region of ovule and arrived at the filiform apparatus of the receptive synergid cell. During this timeframe, three quarters to half of the NTA-GFP signal moved to the micropylar end of synergid cells, indicating that the approach of the pollen tube triggers NTA-GFP accumulation at the filiform apparatus. As reported in [12] and [10], the arriving pollen tubes paused their growth outside the filiform apparatus for 30–50 min, presumably for cell-to-cell communication. During this period, the NTA-GFP signal continued to shift toward the filiform apparatus. At 70–80 min after pollen tube arrival at the micropyle, NTA-GFP signal was only detected at the filiform apparatus and pollen tubes resumed growth and ruptured to release the sperm cells (Figures 1C, D and Movies S1 and S2). Even though only one of the synergids receives the pollen tube, NTA-GFP accumulated at the filiform apparatus in both synergid cells in response to pollen tube arrival, similar to the activation of [Ca^2%^]_cyto_ oscillations in both synergids during pollen tube reception reported in [12].

### Golgi do not concentrate at the filiform apparatus during pollen tube reception

We previously determined that NTA is sequestered in a Golgi-associated compartment in synergid cells that have not attracted a pollen tube [21]. Our live-imaging data suggest that NTA-GFP is selectively moved out of the Golgi and transported to the filiform apparatus in response to pollen tube arrival. However, it is possible that the observed NTA-GFP accumulation at the filiform apparatus is a result of massive reorganization of subcellular compartments. To distinguish between these possibilities, we investigated the behavior of the Golgi in synergid cells during pollen tube reception. We used the semi-*in vivo* imaging system described above with a synergid-expressed Golgi marker (Man49-mCherry) co-expressed with NTA-GFP [33]. In all replicates, the Golgi marker was distributed throughout the synergids, excluded from the filiform apparatus, and co-localized with NTA-GFP as reported previously (Figure 2A, n=93). When a pollen tube approached the synergids, NTA-GFP accumulated at the filiform apparatus of the synergids as observed previously (Figure 1), but the Golgi-mCherry marker remained consistently distributed throughout the synergid cells and did not concentrate near the filiform apparatus (Figure 2B). In order to examine the behavior of the Golgi during later stages of pollen tube reception, we used the synergid-expressed Golgi marker line (*LRE*_*pro*_*::Man49-mCherry*) and pollen that was expressing GFP (*Lat52*_*pro*_*::GFP*). In all cases, the Golgi marker remained randomly distributed throughout the synergid cells, even after pollen tube rupture (Figures 2C-D, Movies S3 and S4, n=106). These results indicate that the accumulation of NTA-GFP at the filiform apparatus during pollen tube reception is not linked to mass redistribution of Golgi.

**Figure 2.**
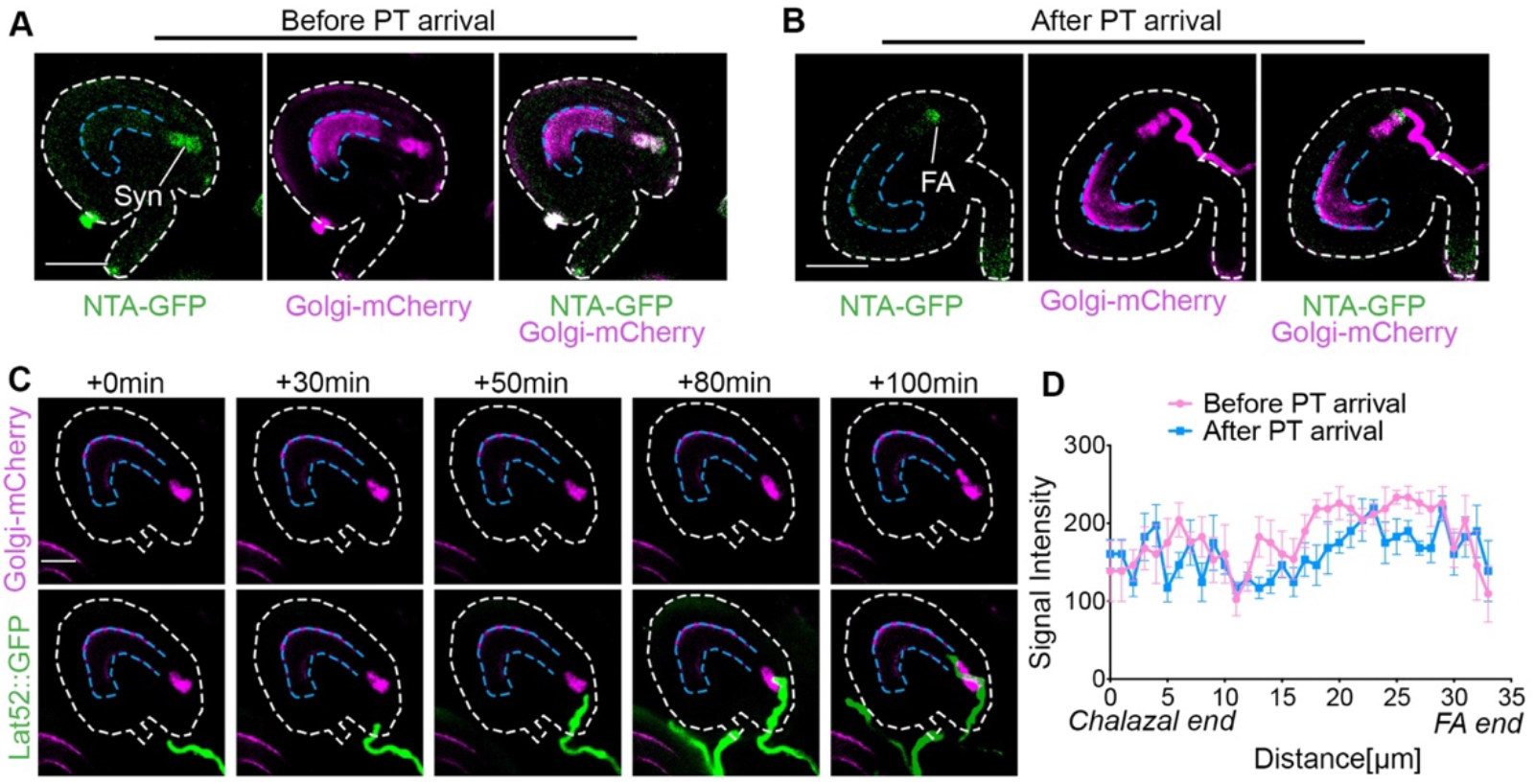
A Golgi marker is randomly distributed throughout synergids during pollen tube reception. (A) NTA-GFP (green signal) and Golgi-mCherry signals (magenta signal) are evenly distributed along the length of the synergid and co-localized within synergid cells before pollen tube arrival. (B) After pollen tube arrival, NTA-GFP accumulated at the FA, but Golgi-mCherry did not accumulate at the FA. (C) Live imaging of Golgi-mCherry during reception of *Lat52::GFP* labeled pollen tubes. (D) Quantification of Golgi-mCherry signal along the length of synergids shown in (C) before and after pollen tube arrival. Bars=30μm (A-B), 50μm (C).

### Endosomes accumulate at the filiform apparatus during pollen tube reception

A signal from the arriving pollen tube seems to trigger the movement of NTA-GFP out of the Golgi-associated compartments. It is possible that pollen tube arrival triggers other changes to synergid subcellular organization. We previously reported the localization of synergid-expressed markers for the ER, peroxisome, endosome and the trans-Golgi Network (TGN) in unfertilized ovules *in vivo* using confocal laser scanning microscopy [33]. Before pollen tube arrival, SP-mCherry-HDEL (an ER-associated marker), mCherry-SKL (a peroxisome-associated marker), and mCherry-RabA1g (a recycling endosome-associated marker) were all distributed evenly throughout synergid cells (Figures S2-S4 and 3A; [33]. The TGN-associated marker SYP61 exhibited two types of distribution patterns before pollen tube arrival: in type 1 synergids, the marker accumulated near the filiform apparatus, whereas type 2 synergids displayed a punctate distribution pattern throughout the cells (Figure S3; [33]. During pollen tube reception, no change was seen in the TGN marker distribution in either type of synergids (Figure S3, Movies S5 and S6). Likewise, the ER and peroxisome markers maintained a diffuse distribution throughout the synergids and did not accumulate at the filiform apparatus (Figures S2 and S4, Movies S7-10). In contrast, we detected a more dynamic behavior of the endosome marker during pollen tube reception. Endosomes are membrane-bound compartments that are involved in the endocytic membrane transport pathway from the plasma membrane to the vacuole. Endosomes also transport molecules from the Golgi and either continue to the vacuole or recycle back to the Golgi [34]. We previously reported that mCherry-RabA1g is distributed throughout synergid cells and had some overlap with NTA-GFP in synergids of unpollinated ovules [33]. Using the semi-*in vivo* system, we confirmed that before pollen tube arrival, mCherry-RabA1g distributed throughout synergid cells (Figure 3A). Interestingly, as pollen tubes approached, the endosome marker started to accumulate at the filiform apparatus of the synergid cells (Figures 3A and B). By the time pollen tube reception was completed, most of the endosome signal was concentrated at or near the filiform apparatus (Figures 3B-F and S5, Movies S11 and S12). These results indicate that the RabA1g endosome compartments have a distinct response to pollen tube arrival and may play a role in facilitating the intercellular signaling pathway that occurs between the synergids and the pollen tube.

**Figure 3.**
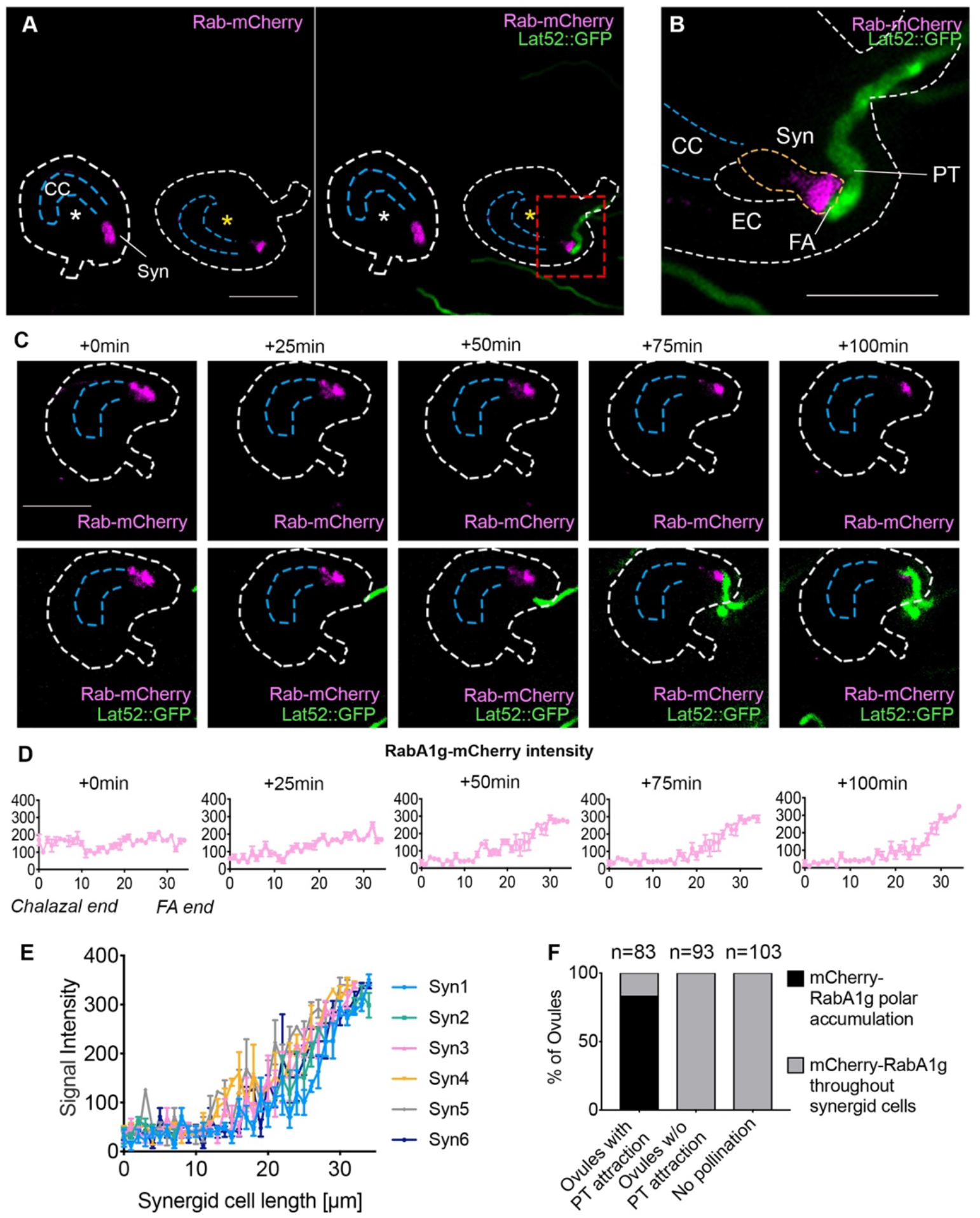
RabA1g endosomes polarly accumulate toward the filiform apparatus during pollen tube reception. (A) RabA1g-mCherry endosome marker (magenta signal) accumulates at the FA region in response to pollen tube arrival (ovule with yellow star). The ovule with no pollen tube attraction (white star) serves as a negative control imaged under the same conditions. (B) Higher magnification of the micropylar region of starred ovule in (red box in panel A). (C) Timing of RabA1g-mCherry polar accumulation during pollen tube arrival. Bars=50μm. (D) Quantification of RabA1g-mCherry signal along the length of synergids during pollen tube reception. Synergid cell from chalazal end to filiform apparatus (FA) end was defined from 0 to 33 μm in length. (E) 6 more examples of the quantification of RabA1g-mCherry signal after pollen tube arrival. (F) Quantification of ovule percentage with RabA1g endosome marker throughout the synergids (gray bars) or with polar accumulation at or near the filiform apparatus (black bars).

### Pollen tube-independent targeting of NTA to the filiform apparatus is not toxic to the synergids

The selective targeting of NTA-GFP from the Golgi to the filiform apparatus during pollen tube arrival (Figures 1 and 2) suggests that NTA trafficking to the pollen tube/synergid interface is important for the intercellular communication process that occurs between the pollen tube and synergids. In *nta-1* mutants, around 30% of ovules display pollen tube overgrowth and fail to complete double fertilization, but the other 70% are fertilized normally [13]. This indicates that NTA is not absolutely required for pollen tube reception, but may function as a modifier of the signaling pathway. Since FER signaling in the synergids leads to cell death as pollen tube reception is completed [17, 23, 24], NTA trafficking to the filiform apparatus could be a mechanism to regulate this death and would thus require sequestration of “toxic” NTA in the Golgi before pollen tube arrival. To test this hypothesis, we took advantage of sequence variation leading to differential subcellular localization of MLO proteins to manipulate the subcellular localization of NTA. When expressed in synergids, other proteins from the Arabidopsis MLO family have different subcellular localization patterns, indicating that specific sequences within the MLOs direct them to different parts of the secretory system [35]. MLO1-GFP localizes at the filiform apparatus when it is ectopically expressed under control of the synergid-specific *MYB98* promoter and cannot complement the *nta-1* pollen tube reception phenotype [21], and Figure 4C and 4G). Domain swaps between different regions of NTA and MLO1 revealed that the C-terminal cytoplasmic tail including the CaMBD of MLO1 (NTA-MLO1^CTerm^, see diagram in Figure 4D) was sufficient to direct the fusion protein to the filiform apparatus of the synergids, in a pattern very similar to MLO1-GFP (Figure 4C). Quantification of the GFP signal along the length of the synergids from the chalazal end to the filiform apparatus in the NTA-GFP, NTA-MLO1^Cterm^-GFP, and MLO1-GFP confirmed that the MLO1 tail was sufficient to cause NTA protein accumulation at the filiform apparatus (Figure 4A, 4C and 4D). In MLO1-GFP and NTA-MLO1^Cterm^-GFP ovules, the majority of GFP signal was detected in the 20-40% of the micropylar pole of synergids and most of the signal overlapped with the diffuse FM4-64 staining at the filiform apparatus (Figure 4F). In contrast, wild-type NTA-GFP is excluded from the filiform apparatus (Figure 4A and [21].

**Figure 4.**
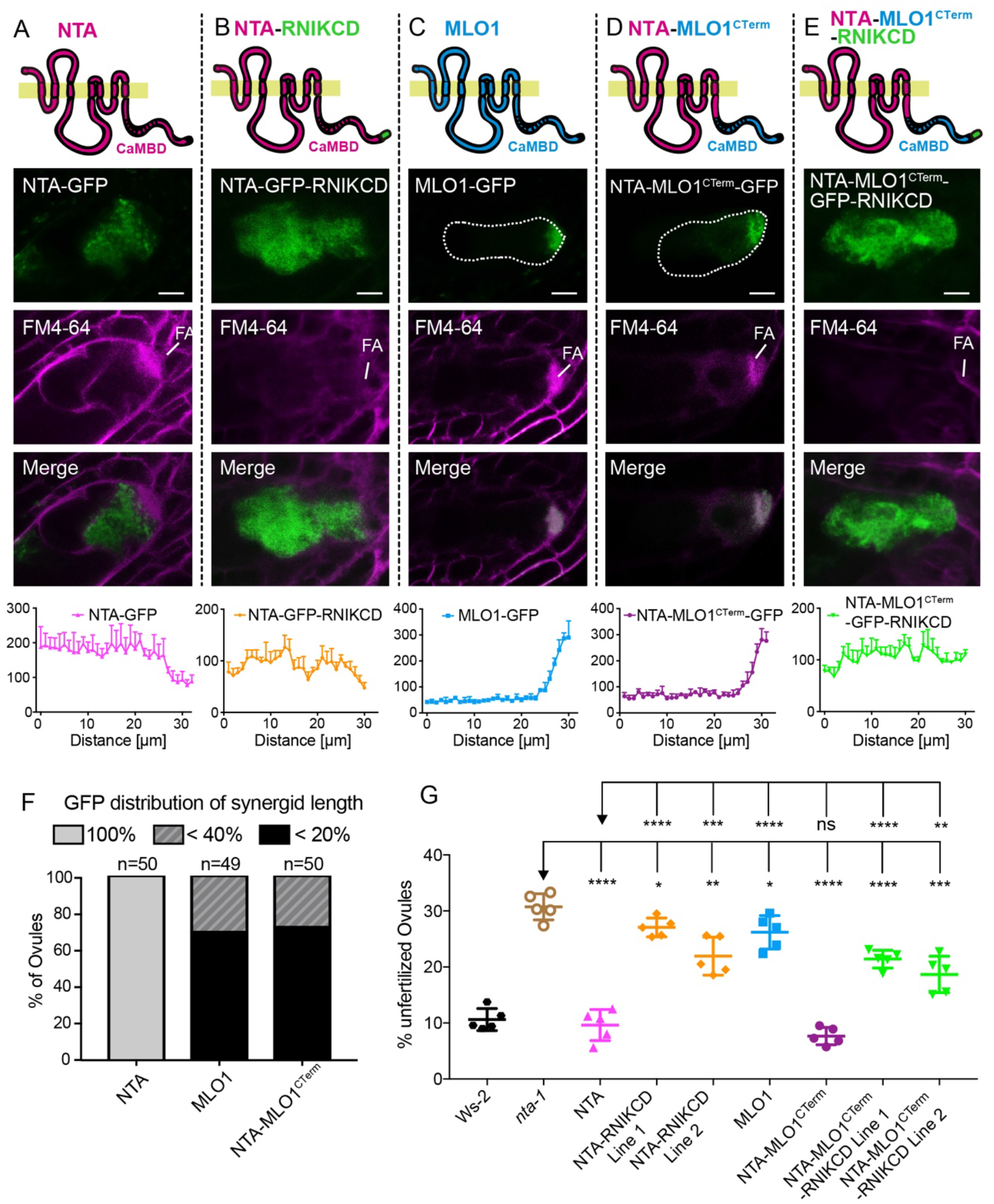
Targeting of NTA to the filiform apparatus before pollen tube arrival is not toxic to synergid cells. (A-E) Localization patterns of MYB98 promoter driven MLO-GFP variants (green signal) in synergids of mature virgin ovules stained with FM4-64 (magenta signal) to reveal the outline of the synergid and the filiform apparatus (FA, diffuse magenta signal). Graphs in each panel show quantification of the GFP intensity of the MLO variants along the length of the synergids. Bars = 10 μm. (F) Percentage of ovules showing MLO-GFP signal throughout the synergids (100% of length), in the FA only (20% of length) and the region surrounding and including the FA (40% of length). (F) Scatter plot of unfertilized ovule percentages in homozygous plants of pMYB98::MLO-GFP in *nta-1* mutants to assess the ability of the MLO-GFP constructs to complement *nta-1*. WS, Wassilewskija. Significance was determined by a Student’s *t*-test (****, *P* < 0.0001; *, *P* = 0.0281; and ns, *P* = 0.2020).

The successful manipulation of NTA subcellular localization provided a tool for determining the functional relevance of NTA redistribution. We transformed the NTA-MLO1^CTerm^-GFP construct into *nta-1* plants and used the percentage of unfertilized ovules as a measurement for the ability of the fusion construct to complement the *nta-1* phenotype of unfertilized ovules caused by pollen tube overgrowth [13]. Synergid-expression of NTA-MLO1^CTerm^-GFP rescued the *nta-1* pollen tube reception phenotype (Figure 4G), indicating that 1) the MLO1^CTerm^ domain is functionally equivalent to the NTA^CTerm^ domain when the protein is localized at the filiform apparatus and 2) premature targeting of NTA to the filiform apparatus is not toxic to synergid cells.

### Polar accumulation of NTA at the filiform apparatus is necessary for pollen tube reception

Early targeting of NTA-MLO1^CTerm^-GFP to the filiform apparatus was sufficient to complement the *nta-1* pollen tube reception phenotype, but did not give an indication of whether filiform apparatus targeting is necessary for NTA function. Since NTA accumulation is triggered by the arrival of a growing pollen tube, drugs that block membrane protein trafficking are not appropriate to inhibit NTA accumulation at the filiform apparatus because they would also inhibit pollen tube growth. Therefore, we used a genetic approach to inhibit NTA trafficking to the filiform apparatus by adding a Golgi retention signal (RNIKCD) to the C-terminus of our MLO-GFP variants. This Golgi retention signal has been reported to cause Golgi-retention when added to proteins that would normally be targeted to the trans-Golgi network and tonoplast [36].

To test if RNIKCD works on MLO proteins, we added RNIKCD to the C-terminus of MLO1 (MLO1-RNIKCD) and NTA (NTA-RNIKCD) and co-infiltrated tobacco leaves with these constructs and the Golgi-mCherry marker (*35S::Man49-mCherry*). Both NTA and NTA-RNIKCD co-localize with Golgi-mCherry marker in tobacco leaves (Figure S7A), as was previously reported for NTA-GFP [21]. However, in contrast to MLO1-GFP which accumulates at the plasma membrane and not in the Golgi, most of MLO1-RNIKCD was retained in the Golgi (Figure S7A). We next added RNIKCD to MYB98_pro_-driven NTA and NTA-MLO1^CTerm^ (NTA-RNIKCD and NTA-MLO1^CTerm^-RNIKCD) and transformed the constructs into *nta-1* and Golgi-mCherry plants for functional and subcellular localization studies, respectively. In synergids of emasculated ovules, both NTA-RNIKCD and NTA-MLO1^CTerm^-RNIKCD fusion proteins co-localized with the Golgi marker (Figure 4B, 4E and S7B), indicating that MLO fusion proteins were mostly retained in the Golgi. Synergid-expressed NTA-RNIKCD and NTA-MLO1^CTerm^-RNIKCD could only partially rescue the *nta-1* pollen tube reception phenotype (Figure 4G), indicating that filiform apparatus-localized NTA is both necessary and sufficient for pollen tube reception.

### Disrupting the NTA CaMBD compromises NTA’s ability to accumulate at the filiform apparatus during pollen tube reception

The dramatic differences in protein localization between NTA-GFP and NTA-MLO1^CTerm^-GFP revealed that the C-terminal cytoplasmic domain of NTA contains sequences that cause NTA to be sequestered in the Golgi before pollen tube arrival and possibly also to allow for directed transport to the filiform apparatus in response to signals from the arriving pollen tube. The C-terminal cytoplasmic domains of MLO proteins contain a predicted CaMBD followed by an unconserved tail of variable length [37–39]. A comparison of C-terminal domains (Figure 5A) revealed that NTA and MLO1 share the conserved 20 amino acid 1-8-14 CaM binding motif of hydrophobic residues interspersed with basic residues that define the MLO CaMBD [38], but have different residues surrounding the conserved tryptophan that has been shown to necessary for CaM binding in MLO proteins [38, 39]. In addition, NTA and MLO1 have different length C-terminal tails after the CaMBD (Figure 5A). In order to determine if the C-terminal tail after the CaMBD is necessary for NTA sequestration in the Golgi before pollen tube arrival, we made a GFP fusion with NTA truncated just after the predicted CaMBD (NTA^Δ481^, Figure 5A) expressed under control of the synergid-specific MYB98 promoter. This construct complemented the *nta-1* pollen tube reception phenotype and localized in the Golgi before pollen tube arrival (Figure 5C and 5E), revealing that the C-terminal tail after the CaMBD is not necessary for NTA sequestration or function. We next tested whether disruption of the CaMBD would affect NTA localization and/or function. A point mutation in the conserved tryptophan necessary for CaM-binding in other CaMBDs [38, 40, 41] was introduced into our NTA-GFP expression construct (NTA^W458A^, Figure 5A) and transformed into the *nta-1* background. In contrast to wild-type NTA and NTA^Δ481^, which result in an even higher fertility in *nta-1* mutants than in the Ws controls, NTA^W458A^ only partially complemented the *nta-1* fertility phenotype (Figure 5E). Before pollen tube arrival, NTA^W458A^ co-localizes with a Golgi marker in synergids similar to wild-type NTA (Figure 5D and Figure S6). However, during pollen tube reception, NTA^W458A^ displays different accumulation patterns that correlate with the ability of the pollen tube to rupture. In our semi-*in vivo* system, ovules with normal pollen tube rupture had at least partial NTA^W458A^ accumulation at the filiform apparatus, while ovules with pollen tube overgrowth did not accumulate NTA^W458A^ at the filiform apparatus (Figure 6). These data reveal that an intact CaMBD enhances NTA’s redistribution to the filiform apparatus during pollen tube reception and that NTA accumulation at the filiform apparatus promotes pollen tube rupture, while the C-terminal tail after the CaMBD is dispensable for NTA function.

**Figure 5.**
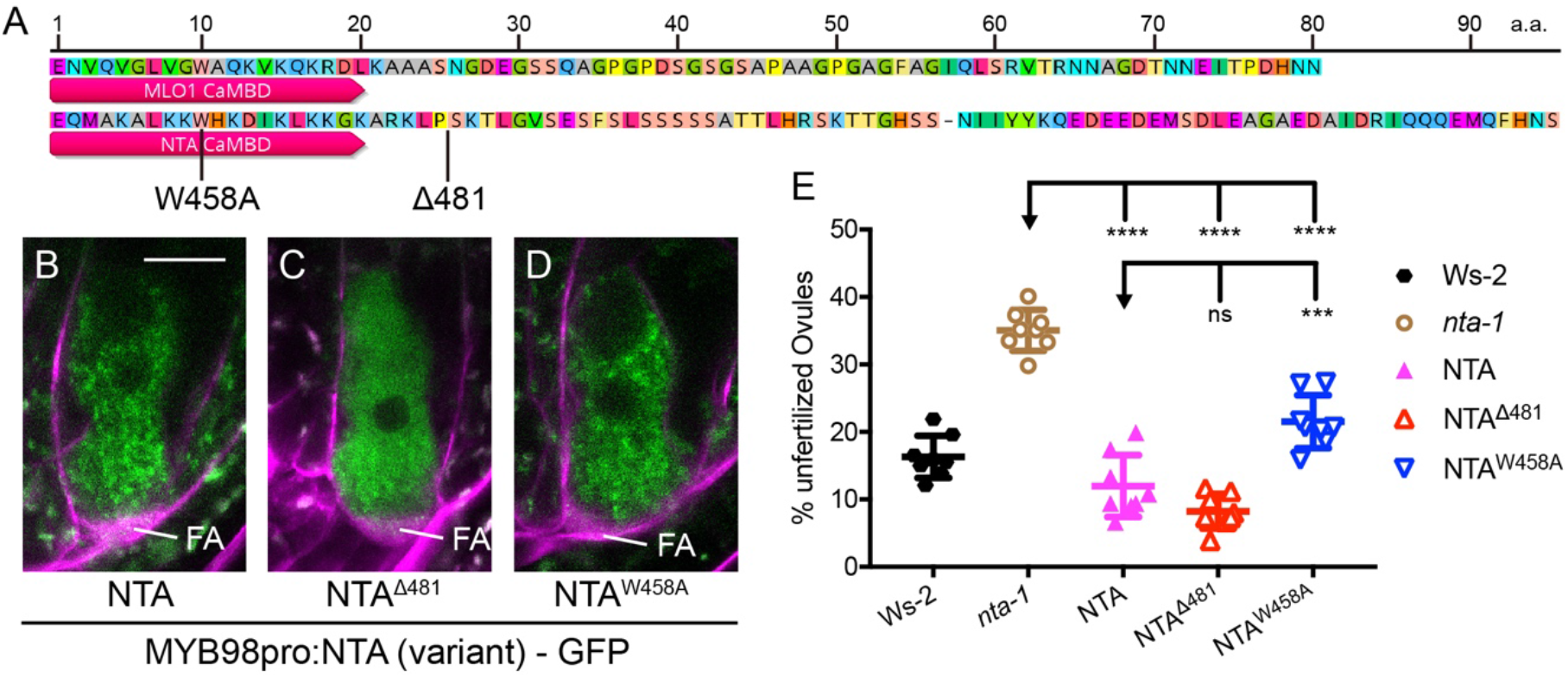
Disruption of the CaMBD affects NTA localization and function. (A) Alignment of MLO1 and NTA C-terminal domains after the seventh membrane span. Red bars highlight the CaMBD; W458A and Δ481 indicate the point mutation in NTA and the deletion point, respectively; a.a., amino acid. (B-D) NTA (variant)-GFP (green) distribution in synergid cells of unfertilized ovules stained with FM4-64 (magenta signal). Bar = 10 μm. (E) Complementation analysis of NTA variants in T2 plants homozygous for MYB98_pro_::NTA(variant)-GFP constructs in *nta-1* mutants. Adjusted P values from a Student’s *t*-test are as follows: **** indicates *P* < 0.0001; *** indicates *P* = 0.001 to 0.0001; and ns indicates *P* > 0.05.

**Figure 6.**
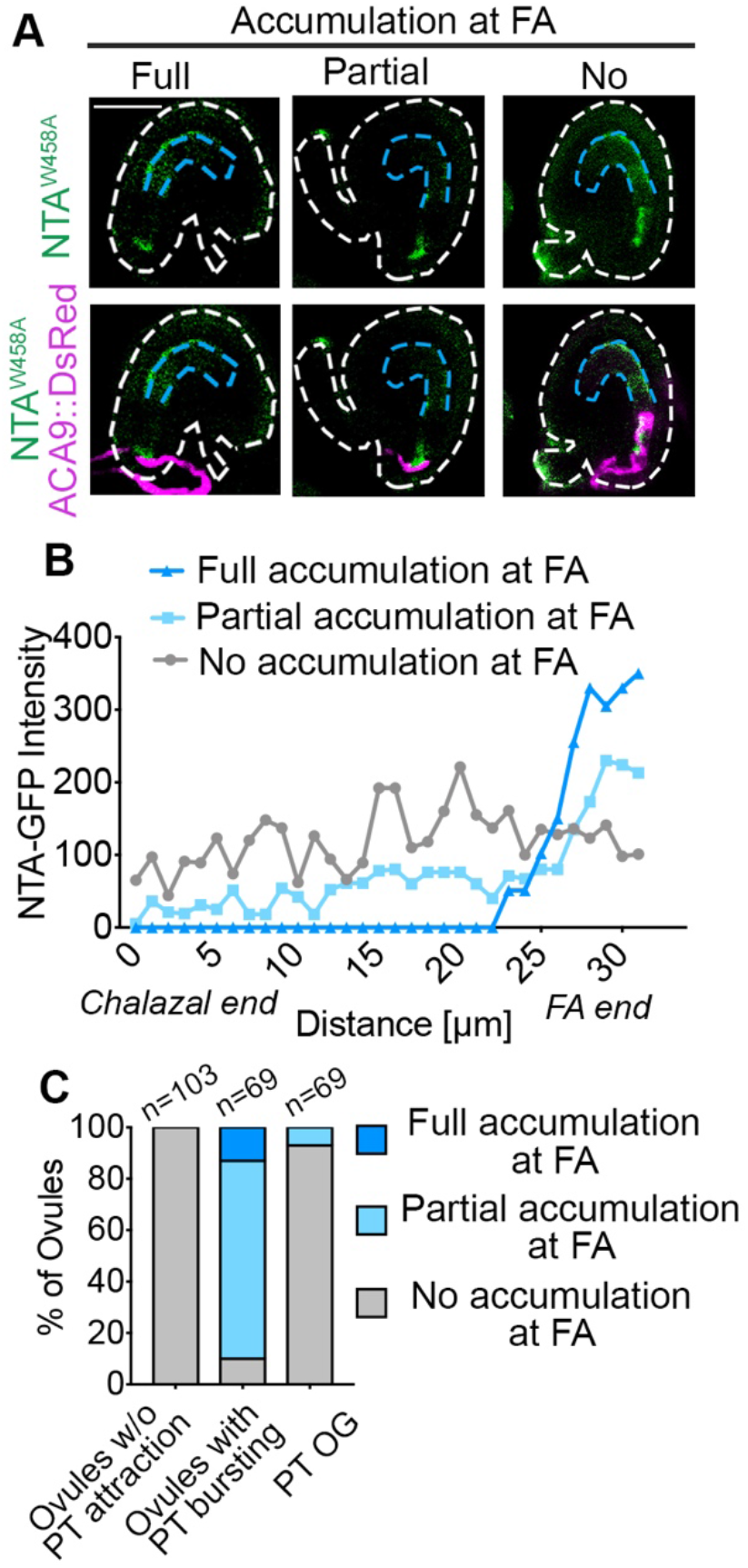
A point mutation in the CaMBD (NTA^W458A^) affects filiform apparatus accumulation of NTA and pollen tube reception. (A) NTA^W458A^-GFP has 3 different localization patterns in response to PT arrival under semi *in-vivo* conditions. Bar=50 μm (B) Quantification of GFP signal intensity in NTA^W458A^ synergids during pollen tube reception. (C) Analysis of NTA^W458A^-GFP distribution patterns in ovules with successful (PT bursting) and unsuccessful (no PT or PT overgrowth (PT OG)) pollen tube reception.

The influence of the CaMBD on NTA accumulation at the filiform apparatus could indicate a connection with the calcium oscillations that occur in synergids during signaling with the arriving pollen tube. The receptor-like kinase FER and the GPI-anchored protein LRE are filiform-localized proteins that interact with each other and form a co-receptor for signals from the arriving pollen tube [14, 16, 17]. *fer* and *lre* mutant ovules fail to initiate synergid calcium oscillations at pollen tube arrival and have highly penetrant pollen tube overgrowth phenotypes [12]. In *nta-1* mutant ovules, synergids have dampened calcium oscillations and around 30% of ovules carrying the mutation display pollen tube overgrowth [12, 13]. In both *fer-1* and *lre-7* backgrounds, NTA-GFP does not accumulate at the filiform apparatus in response to signals from the arriving pollen tube (Figure S8 and [13]), placing NTA accumulation at the filiform apparatus downstream of both FER and LRE and the calcium oscillations conditioned by these proteins.

## Discussion

### Synergids respond to a signal from the approaching pollen tube

Successful pollination and production of seeds requires a series of signaling events between the male gametophyte (pollen tube) and both sporophytic and gametophytic cells of the female. In this study, we used live imaging to characterize dynamic subcellular changes that occur in the synergid cells of the female gametophyte in response to the arrival of the pollen tube. We showed that both the NTA protein and endosomes are actively mobilized to the filiform apparatus where male-female communication occurs during pollen tube reception (Figure 7). Disrupting the ability of NTA to pass through the secretory system by adding a Golgi retention signal compromised its ability to participate in pollen tube reception, revealing that NTA accumulation at the filiform apparatus is necessary for its function. Mutation of NTA’s CaMBD partially compromised NTA redistribution and function in pollen tube reception, revealing that Ca^2%^ may play a role in the synergid response to the signal from the pollen tube.

**Figure 7.**
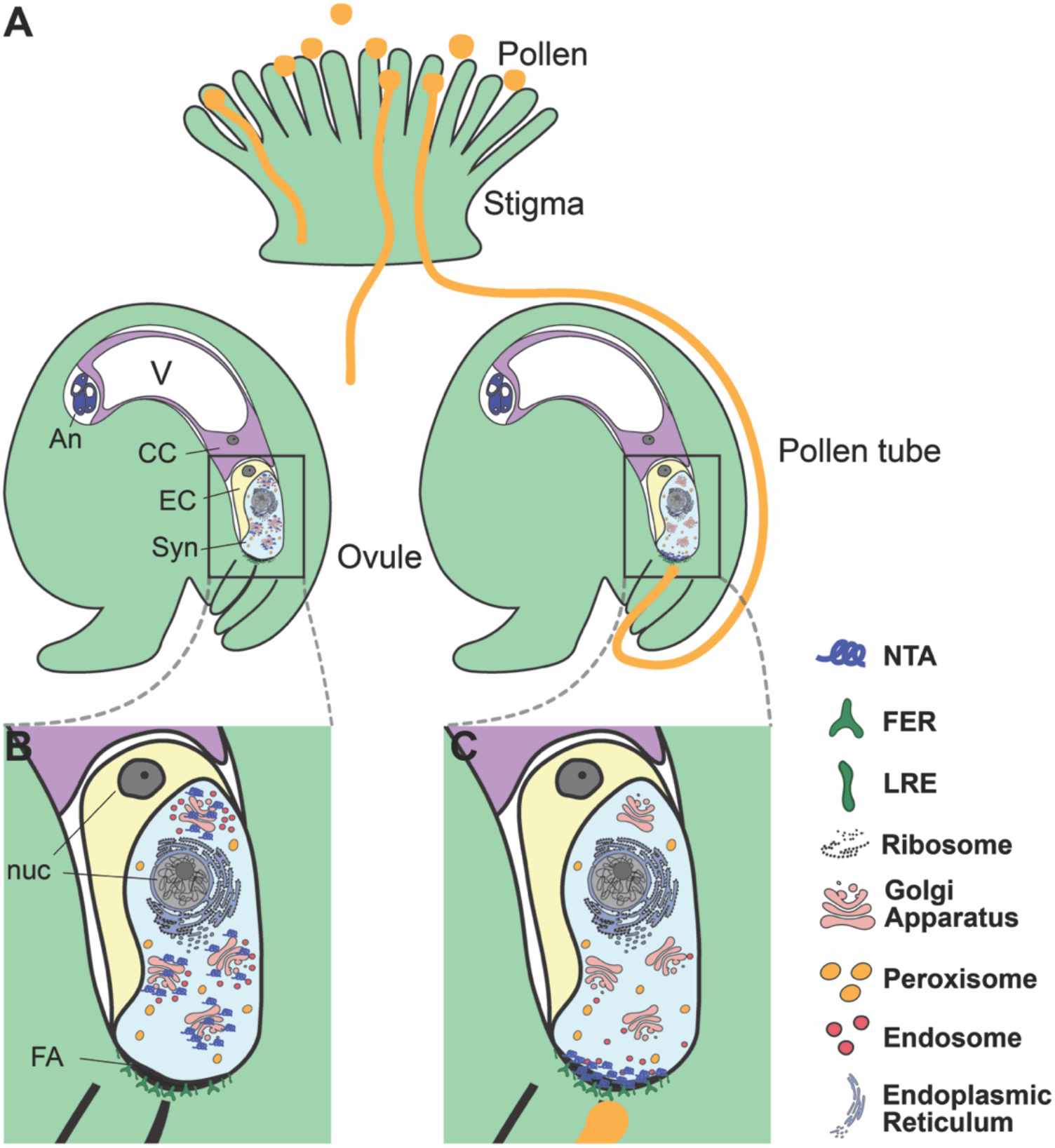
Subcellular dynamics in synergids during pollen tube reception. (A) Pollination using a semi-*in vivo* pollen tube guidance assay. (B) Before pollen tube arrival, NTA is in a Golgi-associated compartment and RabA1g endosomes are distributed throughout the synergids. (C) As a pollen tube arrives, NTA and RabA1g endosomes move toward the filiform apparatus (FA). NTA accumulation is dependent on signaling from the FER receptor like kinase, which acts in a complex with LRE. Abbreviations: CC, Central Cell; Syn, Synergid cells; EC, Egg Cell; An, Antipodal cells; nuc, Nucleus; FA, Filiform Apparatus.

The polar accumulation of the RabA1g endosome marker, but not a Golgi marker (Man49-GFP), near the filiform apparatus during pollen tube reception suggests a change in synergid secretory system behavior that is triggered by the approaching pollen tube. Our results with the ER, Golgi, TGN, and peroxisome markers indicate that the mobilization of the RabA1g endosomes toward the approaching pollen tube is not just a symptom of FER signaling triggering synergid cell death that leads to mass disruption of subcellular compartments. Trans-Golgi Network/Early endosomes (TGN/EEs) have been shown to be involved in the trafficking of both secretory and endocytic cargo [42]. RabA1g is present in endosomes that are highly sensitive to Brefeldin A in roots, suggesting that they could function as recycling endosomes [43]. While the resolution of our live imaging system did not allow us to determine whether NTA completely co-localizes with this compartment, it is tempting to speculate that the RabA1g endosomes are mediating the polar movement of NTA to the filiform apparatus. Alternatively, these endosomes could be transporting other signaling molecules either to or from the filiform apparatus.

### Signal-mediated protein trafficking

Signal-mediated regulation of protein trafficking is an elegant mechanism to control the delivery of molecules to the precise location where they are needed for critical signaling events that occur over relatively short time frames. Selective protein targeting similar to NTA movement in response to pollen tube arrival has also been observed during cell-to-cell communication between the egg and sperm cells in Arabidopsis. After pollen tube reception and release of the sperm cells, a signal from the sperm and/or the degenerated synergid cell causes the egg cell to secrete EGG CELL 1 (EC1) peptides that have been stored in punctate compartments in the egg cytoplasm toward the sperm cells. The sperm cells perceive the EC1 signal and, in turn, mobilize the gamete fusogen HAPLESS2/GENERATIVE CELL SPECIFIC1 (HAP2/GCS1) from a cytoplasmic compartment to the cell surface [44]. This controlled movement of proteins that have already been translated and stored facilitates a quick response to activate the egg and sperm for fertilization. Likewise, NTA mobilization to the filiform apparatus of the synergids as the pollen tube arrives could play a role in sending a signal to the pollen tube that leads to the mobilization of pollen tube proteins that allow the pollen tube to rupture and release the sperm cells. For example, proteins that regulate the integrity of the tip of the pollen tube could be quickly delivered after the “arrival” signal from the synergid is perceived. Recent work on the role of the pollen-expressed ANXUR1 and 2 and BUDDHA PAPER SEAL1 and 2 receptor-like kinases in pollen tube tip integrity support this hypothesis. During pollen tube growth through the female tissues, RALF4 and RALF19 peptides that are secreted from pollen tubes act as ligands for ANX1/2 and BPS1/2 to promote tip growth, while RALF34 secreted from the synergids displaces RALF4 and 19 from the receptors leading to subcellular changes that result in pollen tube rupture [45].

### Calcium and NTA movement

Our study revealed that an intact CaMBD facilitates the trafficking of NTA from the Golgi to the filiform apparatus in response to a stimulus from the approaching pollen tube. Our live imaging studies and previous reports all suggest that the FER/LRE co-receptor is necessary for perceiving this stimulus and that two other CrRLK1L genes, HERK1 and ANJEA are also necessary for NTA accumulation at the filiform apparatus [13, 46]. Polar accumulation of NTA-GFP at the filiform apparatus occurs in a similar time frame to the FER/LRE-dependent Ca^2+^_cyto_ spiking in the synergids during pollen tube reception [10, 12]. Subcellular Ca^2+^ spiking occurs in plant responses to both biotic and abiotic external stimuli. Notably, cytosolic Ca^2+^ oscillations occur during pollen tube-synergid interactions, egg-sperm interactions, and in biotrophic interactions between plant cells and both beneficial and harmful microbes (reviewed in [47]). In most cases, the mechanism for decoding Ca^2+^ spikes into a cellular response is not known, but Ca^2+^-binding proteins such as CaM and CaM-like proteins could play a role in relaying Ca^2+^ signals [48]. In *nta-1* mutants, the cytosolic Ca^2+^ oscillations still occur, but at a lower magnitude than in wild-type synergids, suggesting that NTA could be involved in modulating Ca^2+^ flux [12]. A recent study of the pollen tube-expressed MLO5 and MLO9 genes revealed a link between MLO proteins and recruitment of the calcium channel CNGC18 to control Ca^2+^ flux during directional growth [49]. Several CNGC genes are expressed in synergids and could possibly interact with filiform apparatus-localized NTA to regulate the flow of Ca^2+^ ions into or out of the apoplast near the filiform apparatus.

Calcium has also been linked to regulation of endomembrane trafficking (reviewed in [50]. In animals, CaM plays a role in regulating vesicle tethering and fusion [51], and in plants CaM-like proteins are associated with endosomal populations [52]. Thus, it is possible that the CaMBD in NTA is critical for the precise targeting of NTA in response to pollen tube arrival. Another mechanism that has been proposed for Ca^2+^ regulation of protein targeting is that electrostatic interactions between Ca^2+^ and anionic phospholipids in specific domains of the plasma membrane regulate vesicle fusion and differential recruitment of proteins to these domains [53, 54].

An interesting parallel between powdery mildew infection and pollen tube reception is that in both cases, an MLO protein accumulates at the site of interaction with a tip-growing cell. During powdery mildew infection, active transport of proteins and lipids to penetration site leads to membrane remodeling and establishment of the extrahaustorial membrane (EHM) which separates the plant cytoplasm from the invading fungal hyphae [55]. This specialized membrane is enriched in the anionic phospholipid phosphatidylinositol 4,5 bisphosphate (PI(4,5)P_2_). Depletion of PI(4,5)P_2_ in *phosphatidylinositol 4-phosphate 5-kinase 1/2* double mutants (*pip5k1 pip5k2*) resulted in powdery mildew resistance associated with a failure of MLO2 to accumulate at the EHM in early stages of fungal penetration [56]. Although the composition of the filiform apparatus and its relationship with the arriving pollen tube has not been established, it is possible that similar membrane modification occurs in the filiform apparatus during signaling with the pollen tube and is linked to NTA accumulation at pollen tube arrival.

### Cell death and pollen tube reception

MLOs were first discovered in barley as powdery mildew resistance genes [57]. *mlo* mutants in both monocots and dicots are resistant to powdery mildew infection, indicating that the MLO proteins are required for infection. These mutants also have ectopic cell death, indicating that one function of MLO proteins is to negatively regulate cell death [58]. Pollen tube reception is catastrophic for both the pollen tube and the receptive synergid cell: both cells die as a result of successful male-female signaling and delivery of the male gametes. The timing of synergid degeneration remains under debate, with some studies suggesting that it occurs upon pollen tube arrival at the synergid and others concluding that it occurs concurrently with pollen tube rupture [31, 59, 60]. Our live imaging experiments with both NTA-GFP and the subcellular compartment reporters suggest that the synergid cells are still alive and regulating their secretory systems up to the point of pollen tube rupture. Given the function of the “powdery mildew” members of the MLO gene family in regulating cell death [61], it is possible that one role of NTA is to prevent early degeneration of the synergids. In both pollen tube reception and powdery mildew infection, an MLO protein might be needed at the site of penetration to stabilize the cell and prevent precocious cell death. Our result that premature delivery of NTA (in the NTA-MLO1^CTerm^ domain swap construct) to the filiform apparatus of the cell does not disrupt pollen tube reception is consistent with this hypothesis, since other signaling processes occurring in the synergids during communication with the arriving pollen tube would likely not be triggered by simply moving NTA to the filiform apparatus in the absence of a pollen tube.

In this study, we showed that signals from an approaching pollen tube trigger the movement of NTA out of the Golgi and to the filiform apparatus and that this redistribution is necessary for pollen tube reception. Future work will focus on determining the mechanism through which NTA polarly accumulates at the filiform apparatus and on identifying the signals from the pollen tube that lead to important subcellular changes in the synergids during pollen tube reception.

## Materials and methods

### Plant materials and growth conditions

*Arabidopsis thaliana* lines expressing *NTA*_*pro*_*::NTA-GFP*, *MYB98*_*pro*_*::NTA-GFP*, *MYB98*_*pro*_*::MLO1-GFP*,Golgi-associated marker (Man49-mCherry), ER-associated marker (SP-mCherry-HDEL), TGN-associated marker (SYP61-mCherry), peroxisome-associated marker (Peroxisome-mCherry) and recycling endosome-associated marker (mCherry-RabA1g), were generated and described in previous publications [13, 14, 21, 33]. Pollen tube marker lines, *ACA9::DsRed* [62] and *Lat52::GFP* [28] were generously provided by Dr. Aurelien Boisson-Dernier and Dr. Ravi Palanivelu. *fer-1* seeds were provided by Dr. Ueli Grossniklaus and *lre-7* seeds were provided by Dr. Ravi Palanivelu. Seeds were sterilized and plated on ½-strength Murashige and Skoog (MS) plates. All plates were sealed and stratified at 4°C for two days, and then transferred to the growth chamber (long day conditions, 16 h of light and 8 h of dark at 22°C) for germination and growth. After 5-7 days, seedlings were transplanted to soil. Seeds from transformed lines were sterilized and plated on ½ MS plate with 20 mg/L hygromycin for selection of transgenic seedlings, which were then transplanted to soil and grown in long days.

### Live Imaging of Pollination Using a Semi-*in vivo* Pollen Tube Guidance Assay

The semi-*in vivo* system of pollen tube reception was modified from [28]. Approximately 150 μL of pollen germination media (5 mM KCl, 1 mM MgSO_4_, 0.01% (w/v) H_3_BO_3_, 5 mM CaCl_2_, 20% sucrose, 1.5% agarose, and adjusted pH to 7.5 with KOH) was poured into a Glass Bottom Culture Petri Dish (MatTek Corporation, P35G-1.0-20-C) and spread out using a pipette. Pistils were emasculated and 2 d later were hand pollinated with *ACA9::DsRed* or *Lat52::GFP* pollen and returned to the growth chamber. Approximately 1 h after pollination, pistils were removed from plants and placed on double sided tape on a glass slide. Stigmas were cut using single-sided razor blade and placed on pollen germination media using forceps. 8-10 ovules were arranged around the cut style and the petri dish was returned to the growth chamber. After 4-6 h, pollen tubes grew through the stigma and style and emerged near the ovules. Imaging commenced when the pollen tubes approached ovules. Time-lapse images were acquired at 5 min intervals by spinning disk confocal microscopy using an Andor Revolution XD platform with a Yokogawa CSU-X1-A1 scanner unit mounted on an Olympus IX-83 microscope and a 20X/0.5 NA objective (Olympus). An Andor iXon Ultra 897BV EMCCD camera was used to capture GFP fluorescence (488-nm excitation) and red fluorescent protein (dsRed or mCherry) fluorescence (561-nm excitation).

For each experimental condition, at least 60 ovules were imaged over the time course from pollen tube approaching the ovule to completion of pollen tube reception (pollen tube rupture to release the sperm cells). Neighboring ovules that did not attract pollen tubes were imaged at the same time and served as controls for photoxicity.

### Confocal laser scanning microscopy (CLSM)

CLSM to examine MLO variant subcellular localization was performed on ovules dissected 2 d after emasculation. FM4-64 staining was performed according to the protocol described in [21]. CLSM was done using either a Nikon A1Rsi inverted confocal microscope according to [63] or a Leica SP8 upright confocal microscope according to [21].

### Quantification of fluorescence signal intensity

Two-channel images were adjusted for brightness and contrast using Fiji [64]. Then, they were input to NIS-Elements software (Ver. 5.02) to measure the fluorescence signal intensity. A line that spanned from the chalazal end to the filiform apparatus end of the synergid was used for the signal intensity measurements. For a more accurate representation of the total area of the synergid, signal intensities were recorded along the same length line at five parallel position within the synergid cell. Finally, all the measurement data were output as Excel files. Graphs and statistical analysis were performed with Prism software (www.graphpad.com).

### Video processing

Images were filtered to remove the noise and cropped using Fiji (Version 2.0.0). QuickTime Player (Version 10.5) was used for movie editing and time-lapse analyses.

### Cloning and Generation of Transgenic Lines

PCR amplification with PHUSION High-Fidelity Polymerase (NEB, M0535S) or Q5 High-Fidelity DNA Polymerase (NEB, M0419S) were used to generate the following constructs in this study. Genes were amplified with primers that had attB1 and attB2 sites for recombination via BP reaction into the Gateway-compatible entry vector pDONR207. Full-length NTA entry vectors used in this study was generated as described previously [21]. NTA truncation was amplified using NTA full-length entry vector as a template with forward primer NTA-FattB1 and the reverse primer NTA481-RattB2 (See all primer sequences in Table S1). The NTA^W458A^ point mutation was generated using the same *NTA* template and amplifying two fragments of *NTA* with desired point mutations introduced into the primers: NTA-FattB1 + NTAW458A-R and NTAW458A-F + NTA-RattB2. The two fragments were purified and pasted together with overlaps using a PCR-pasting protocol. The NTA-MLO1^CTerm^ construct was generated using the full-length entry vectors of NTA and MLO1 used in previous study [21] as templates and amplifying two fragments of *NTA* and *MLO1* using the two pairs of primers: NTA-FattB1 +NTA-R19 and MLO1-F + MLO1-RattB2. The two fragments were purified and pasted together with overlaps using a PCR-pasting protocol. The NTA-GFP-RNIKCD construct was generated using the full-length expression vector of MYB98_pro_::NTA-GFP as template. Two separate fragments of NTA and GFP-RNIKCD-1 were amplified with two pairs of primers: NTA-FattB1 + NTA-Gly5-GFP-R and NTA-Gly5-GFP-F + GFP-RNIKCD-RattB2. The two fragments were purified and pasted together with overlaps using a PCR-pasting protocol. The NTA-MLO1^CTerm^-GFP-RNIKCD construct was generated using the full-length expression vector of MYB98_pro_::NTA-MLO1^CTerm^-GFP as template and two fragments of NTA-MLO1^CTerm^ and GFP-RNIKCD-2 were amplified using the two pairs of primers: NTA-FattB1 + MLO1-Gly5-GFP-R and MLO1-Gly5-GFP-F + GFP-RNIKCD-RattB2. The two fragments were purified and pasted together with overlaps using a PCR-pasting protocol. The coding sequence from the truncation and the point mutation and the fusion sequences were fully sequenced in entry vectors. NTA^Δ481^ and NTA^W458A^ entries were then recombined via LR reactions into the pMDC83 backbone with the MYB98_pro_ [65] to drive expression of each NTA variant in synergid cells with a C-terminal GFP fusion and NTA-GFP-RNIKCD and NTA-MLO1^CTerm^-GFP-RNIKCD entries were then recombined via LR reactions into pMDC32 plasmids with the MYB98_pro_ for synergid expression and FER_pro_ for tobacco transient expression.

For *nta-1* complementation assay and co-localization analyses, expression vectors were transformed into the *Agrobacterium tumefaciens* strain GV3101 and transformed into the *nta-1* mutant background or the Col-0 background (for co-localization combinations, Col-0 stably expressing the Golgi or TGN synergid secretory markers was used for transformation) via the floral-dip method [66]. Stable transgenic lines were selected using their respective selections described above. Homozygous T2 or T3 lines were used in *nta-1* complementation assay and co-localization imaging in the synergid was done in a T1 or T2 analysis. The NTA_pro_::NTA-GFP construct described in [13] was introduced into the *lre-7* background by floral dip as described above.

### *nta-1* Complementation Assays

2-4 independent insertion lines for each construct were taken to the T2 generation and screened for homozygosity using fluorescence microscopy to ensure transgene expression in synergids of every ovule. Unfertilized vs. fertilized ovule counts from self-pollinated flowers were assessed in at least five plants of each insertion line and compared to *nta-1*, Wassilewskija (Ws-2; wildtype), and the previously published full-length NTA (MYB98_pro_::NTA-GFP in *nta-1* background, [21]. Ovule counts were statistically analyzed using Prism with significance determined using a Student’s *t*-test. Comparisons of the NTA variants were made to full-length NTA and the *nta-1* mutant.

## Acknowledgements

We thank Patrick Day for technical assistance and Rachel Flynn and Thomas Davis for helpful discussions and comments on the manuscript. This work was supported by funds from NSF IOS-1733865 to SAK, Purdue University Start-up funds to SAK, and a grant from the Oklahoma Center for the Advancement of Science and Technology #PS14-008 to SAK. Spinning disk confocal microscopy in the Staiger laboratory was supported, in part, by an award from the Office of Science at the US Department of Energy, Physical Biosciences Program, under contract number DE-FG02-09ER15526 to CJS.

## Author Contributions

JY, YJ, DSJ, and SAK conceived and designed the experiments. JY, YJ, DSJ, NL, and WZ performed the experiments. JY, YJ, DSJ, and SAK analyzed the data. JY, YJ, DSJ, and SAK wrote the manuscript, and all authors revised and approved the final manuscript.

## Competing Interests

The authors declare no competing interests.

## Supplemental Materials

**Supplementary figure 1.**
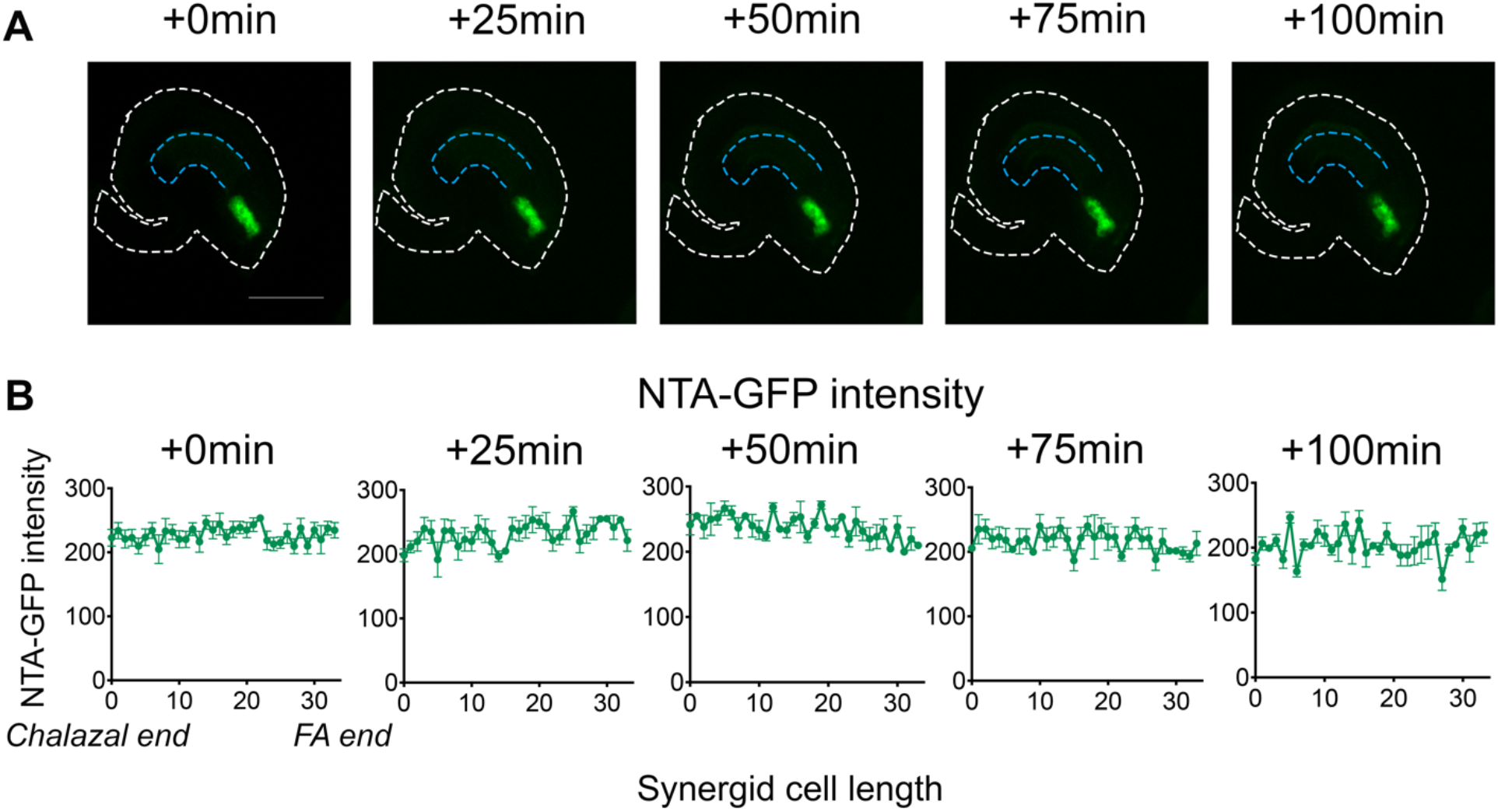
NTA-GFP does not polarly accumulate at the filiform apparatus in synergids without pollen tube attraction. (A) NTA-GFP (green signal) distribution pattern in synergids at 0min, 25min, 50min, 75min, and 100min timepoints, respectively. (B)Signal intensity measurement of NTA-GFP intensity at 0min, 25min, 50min, 75min, and 100min timepoints, respectively. Bar=50μm.

**Supplementary figure 2.**
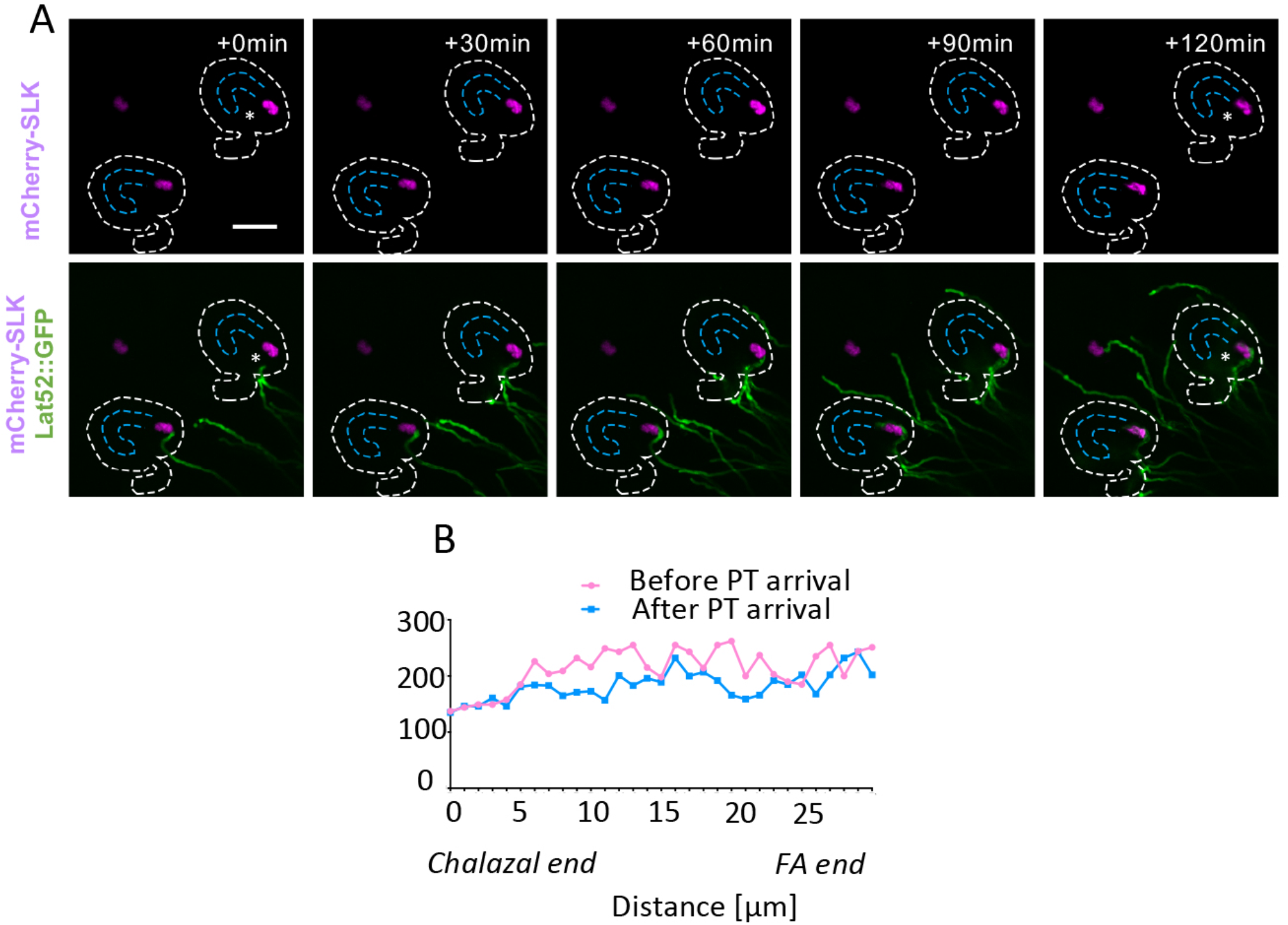
Peroxisomes do not exhibit polar accumulation at the FA during pollen tube reception. (A) The peroxisome marker mCherry-SLK (magenta signal) does not accumulate to the FA region after pollen tube (green signal) arrival. (B) Quantification of mCherry-SLK signal before and after PT arrival in the ovule with the white star. Bars=50μm.

**Supplementary figure 3.**
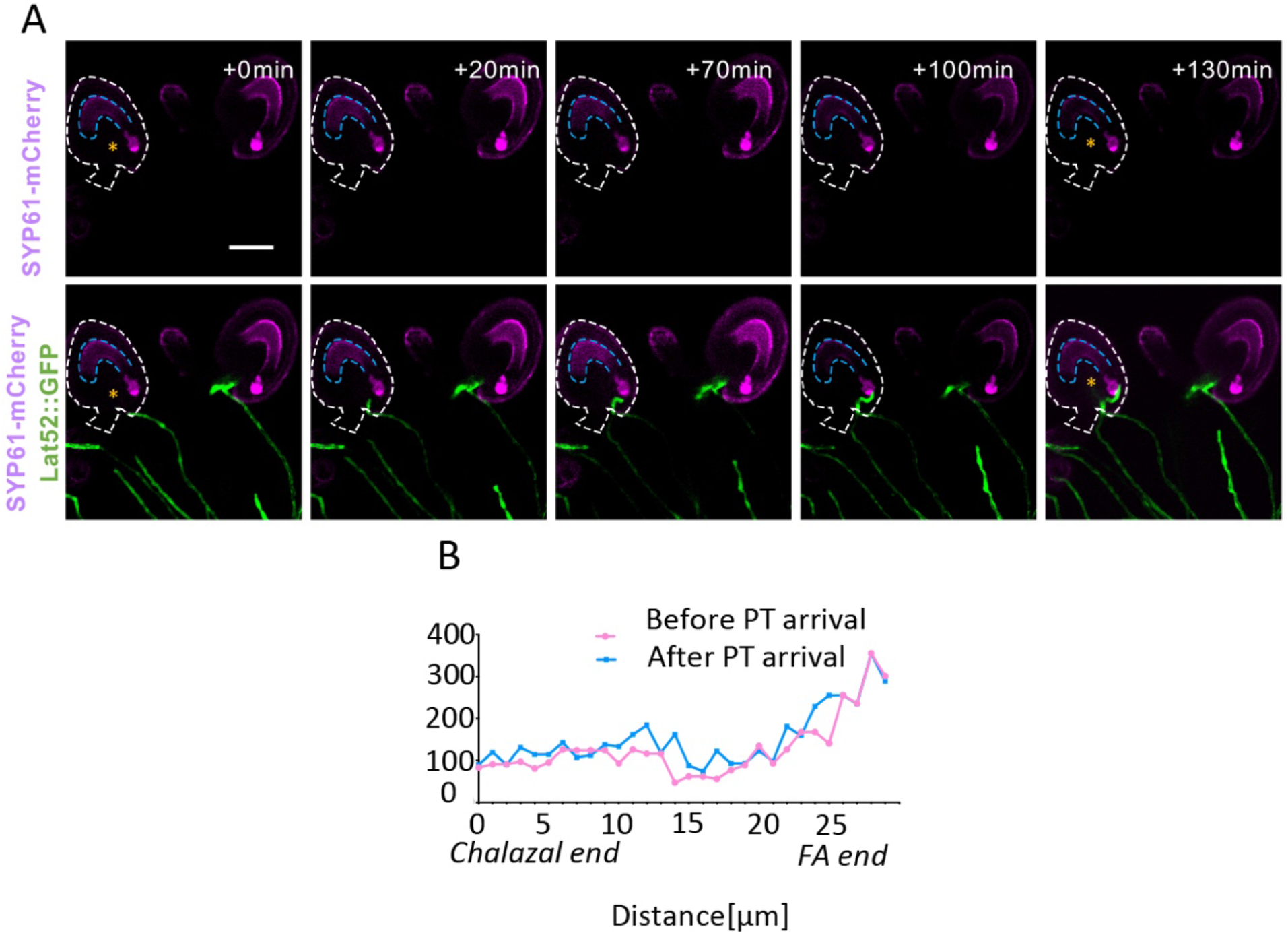
Trans-Golgi marker distribution in synergids does not change in response to pollen tube reception. (A) The trans-Golgi marker SYP61-mCherry (magenta signal) is concentrated toward the micropyle region of synergid cells both before and after pollen tube (green signal) arrival (ovules with yellow stars). (B) Quantification of SYP61-mCherry signal along the length of a synergid before and after pollen tube reception. Bars=50μm.

**Supplementary figure 4.**
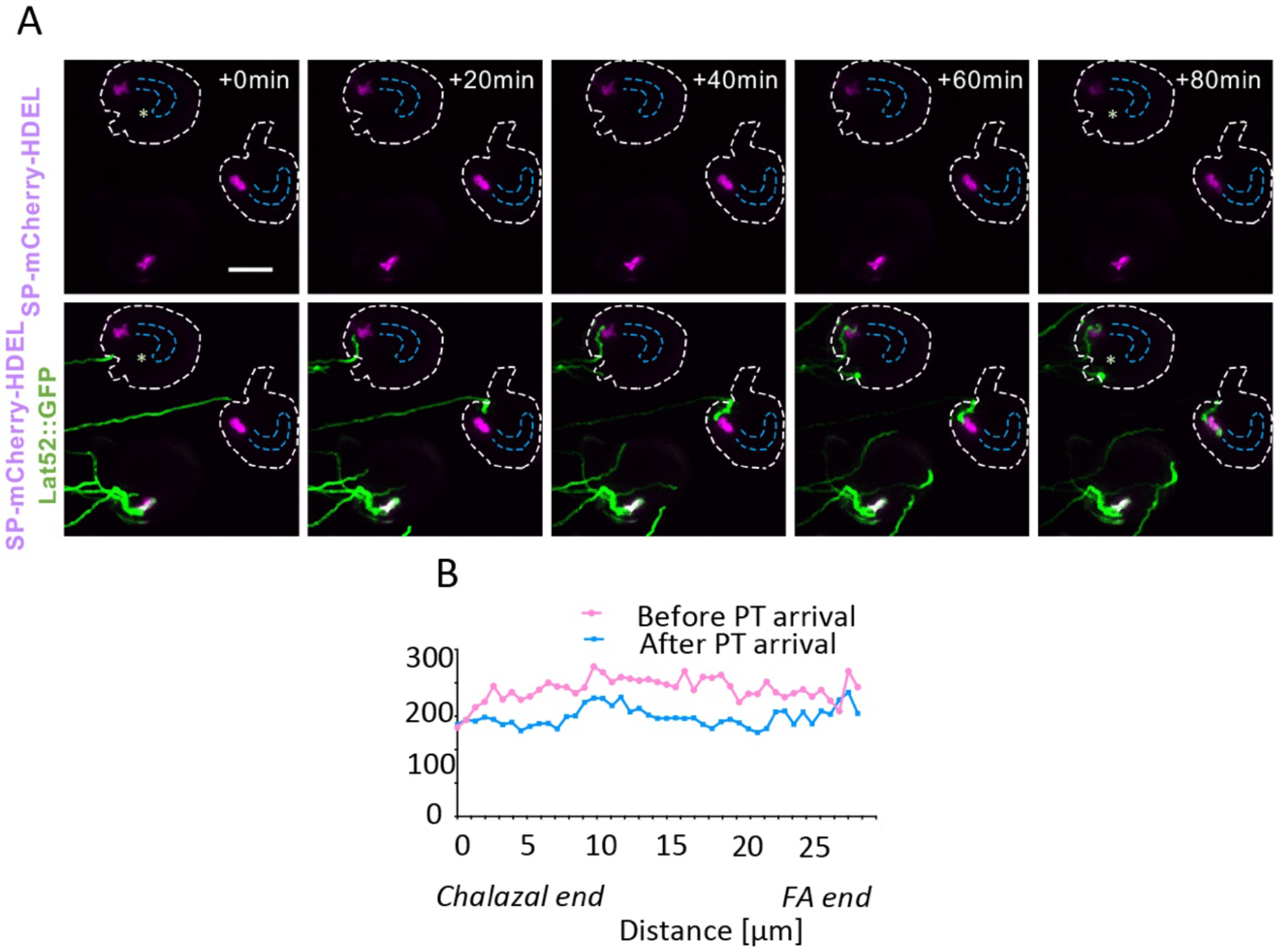
ER marker distribution in synergids does not change in response to pollen tube reception. (A) Before and after pollen tube (green signal) arrival, the ER marker SP-mCherry-HDEL (magenta signal) is distributed throughout synergid cells. (B) Quantification of SP-mCherry-HDEL signal along the length of synergids in ovule with white star before and after pollen tube reception. Bars=50μm.

**Supplementary figure 5.**
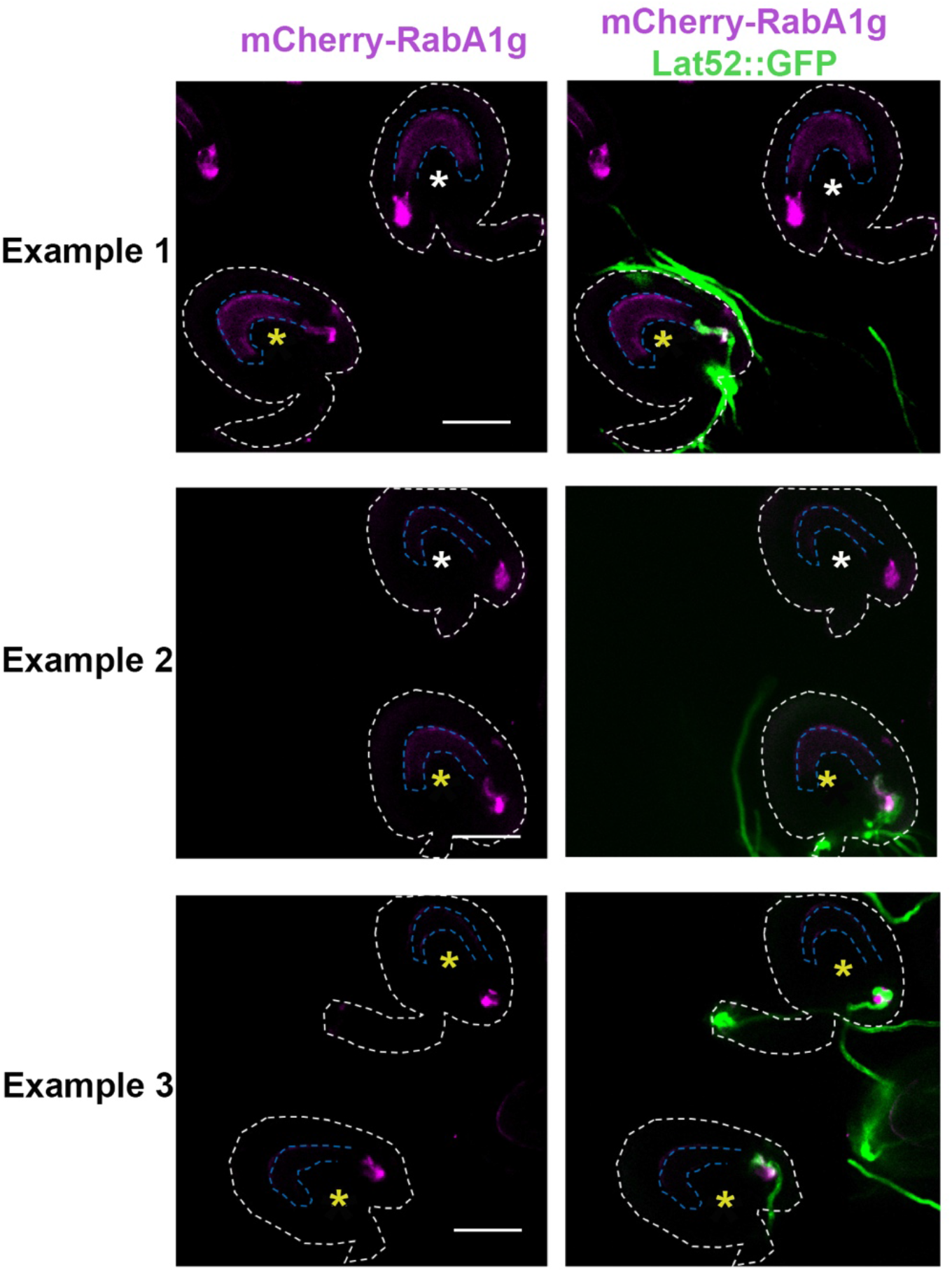
Additional examples of the polar accumulation of RabA1g endosomes during pollen tube reception. Endosome marker accumulates near the filiform apparatus region as pollen tube approaches (ovules with yellow stars), but no polar accumulation was found in ovules that did not attract a pollen tube (ovules with white stars). Bars=50μm.

**Supplementary figure 6.**
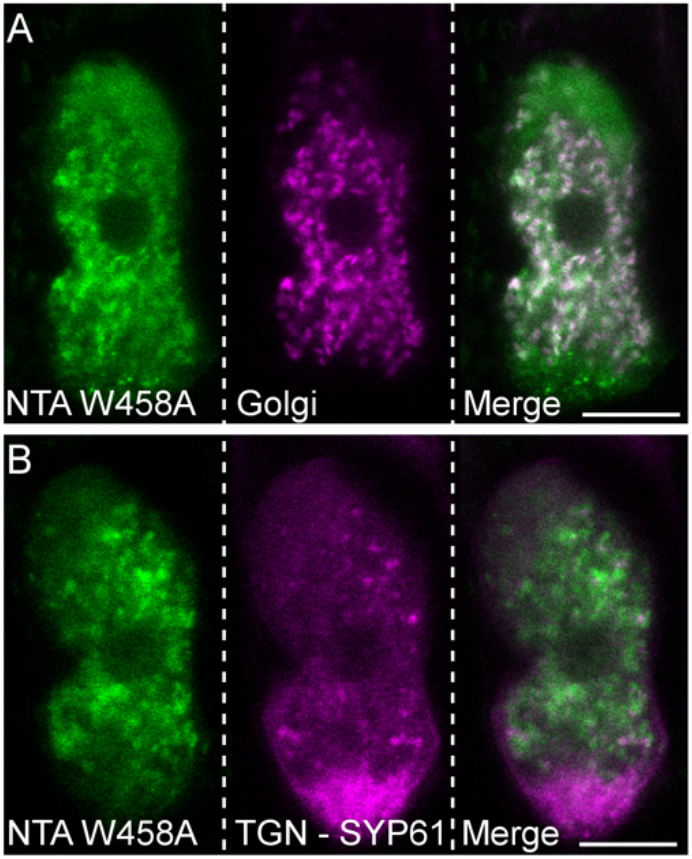
NTA^W458A^ co-localizes with Golgi marker in synergid cells before pollen tube arrival. Colocalization of NTA^W458A^-GFP (green) with Golgi marker (*LRE::proMan49-mCherry*, magenta) (A) and the trans-Golgi network marker (TGN, *MYB98pro::SYP61:mCherry*, magenta) (B) in the synergid cell of unpollinated ovules. Bars = 10 μm.

**Supplementary figure 7.**
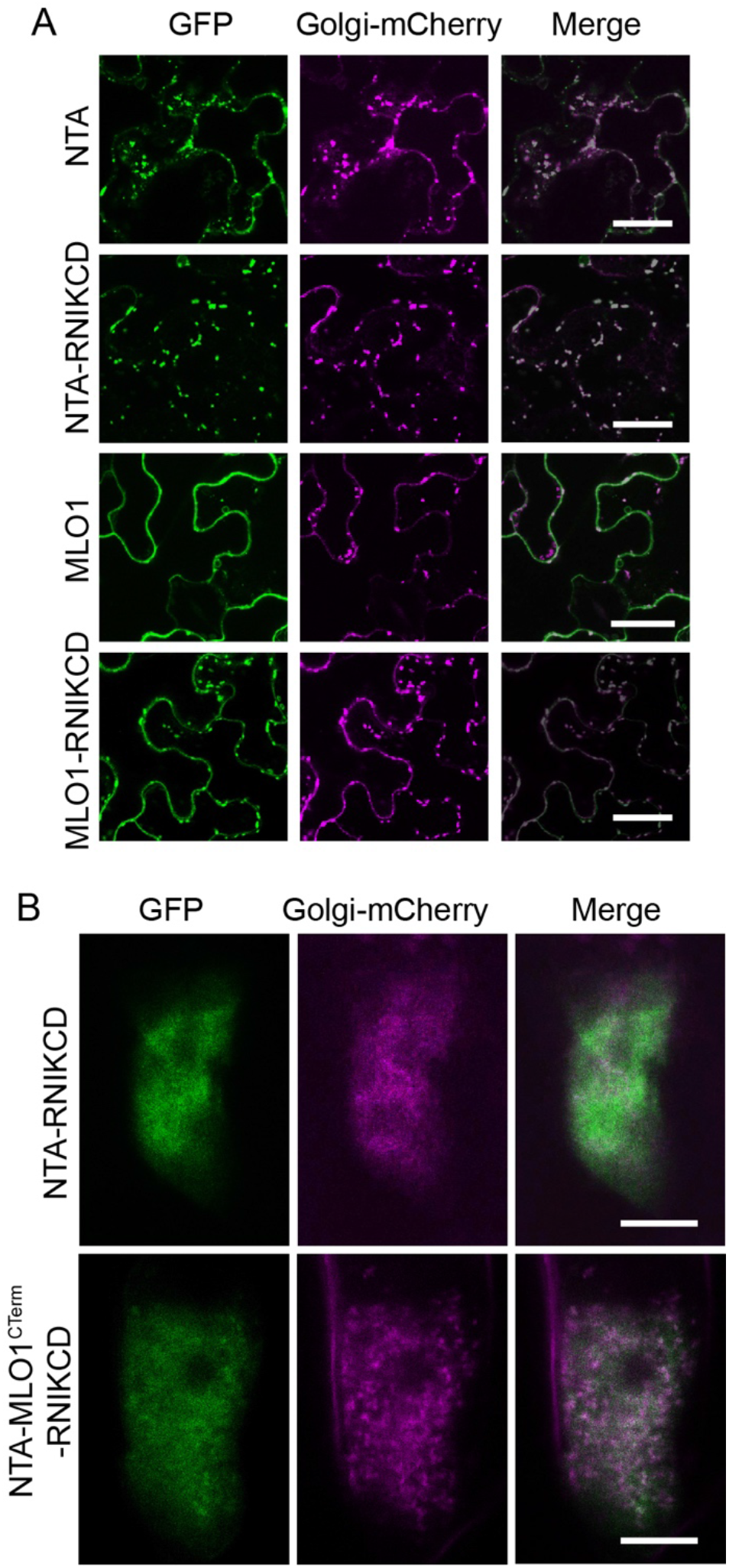
A C-terminal Golgi retention signal retains MLO1 and NTA variants in the Golgi. (A) Colocalization of MLO-GFP (green) with the Golgi-mCherry marker (magenta) indicates that the Golgi retention signal RNIKCD partially retains MLO1 in the Golgi in transiently-transformed tobacco epidermis, Bars = 20 μm. (B) Colocalization of MLO-RNIKCD (green) with the Golgi-mCherry marker (magenta) indicates that the Golgi retention signal RNIKCD retains NTA-MLO1^CTerm^ in the Golgi in the synergid. Bars = 10 μm.

**Supplementary figure 8.**
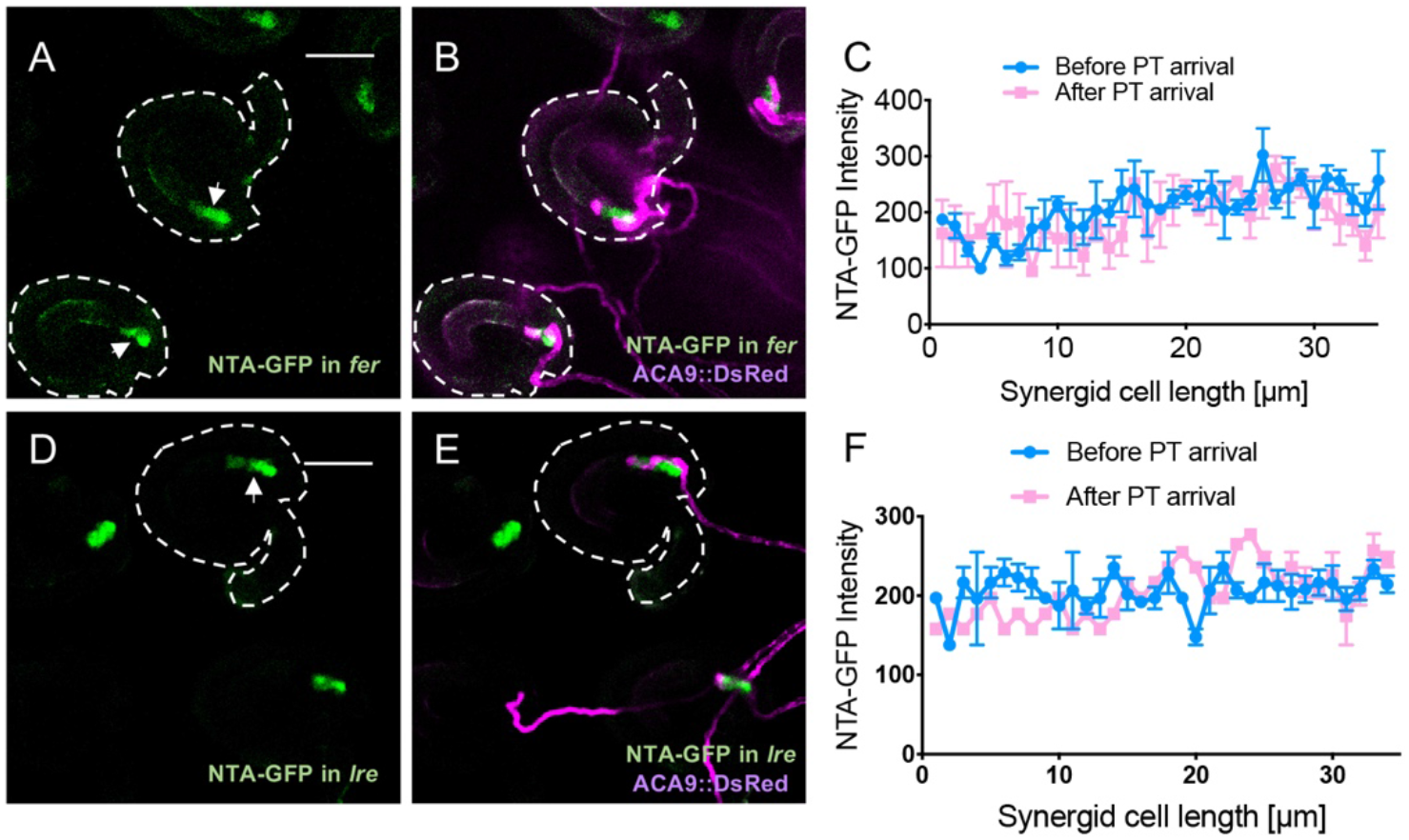
NTA-GFP polar accumulation requires FER and LRE. (A-B) In *fer* mutant synergids, NTA-GFP (green signal) distributes throughout synergids (arrows) with ACA9::DsRed pollen tube overgrowth (magenta signal). (C) Quantification of NTA-GFP signal along the length of *fer-1* synergids before and after pollen tube reception. (D-E) In *lre-7* ovules, NTA-GFP distributes throughout synergids (white arrow) with pollen tube overgrowth. (F) Quantification of NTA-GFP signal along the length of *lre-7* synergids before and after pollen tube reception. FA, Filiform Apparatus. Bars=50μm.

**Supplementary table 1.**
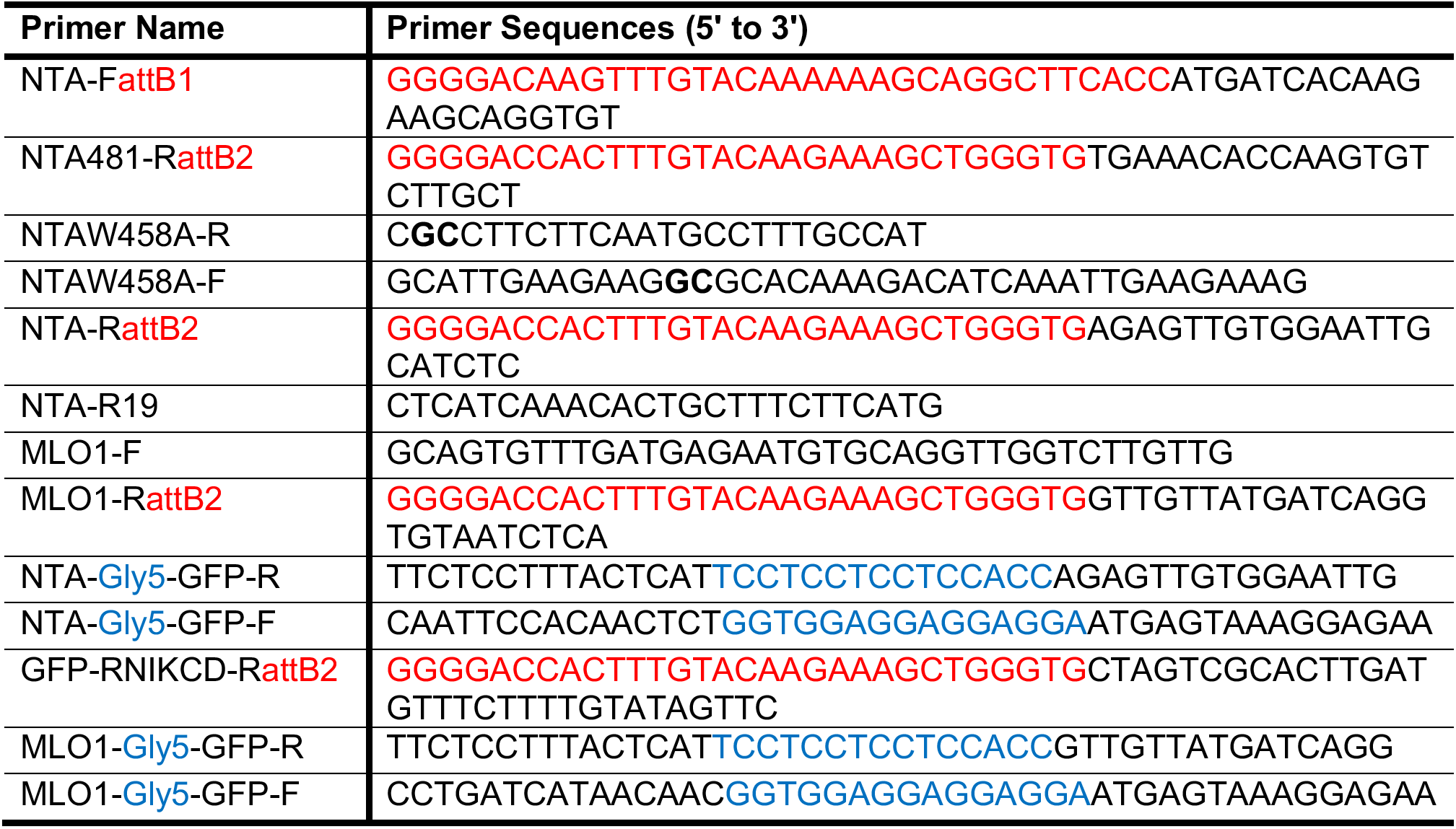
List of primers used for cloning.

## Supplemental Movies

**Supplementary movie 1.**
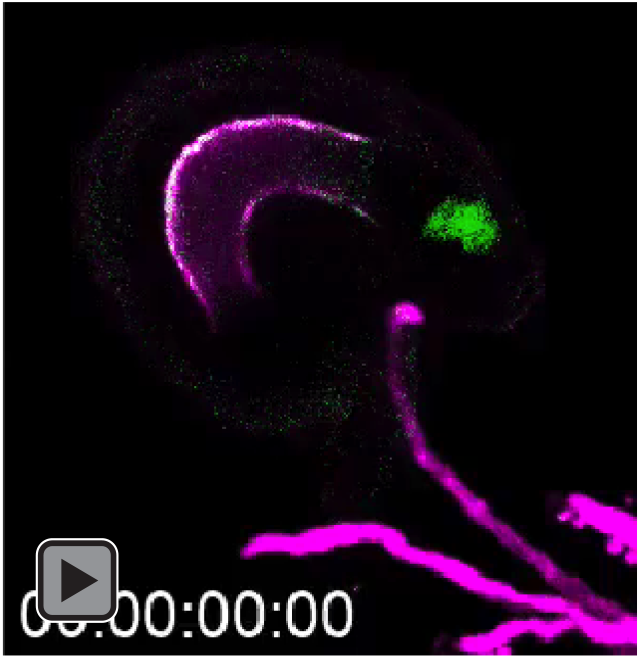
NTA-GFP (green signal) accumulates at the filiform apparatus as ACA9::DsRed labeled pollen tube (magenta signal) approaches.

**Supplementary movie 2.**
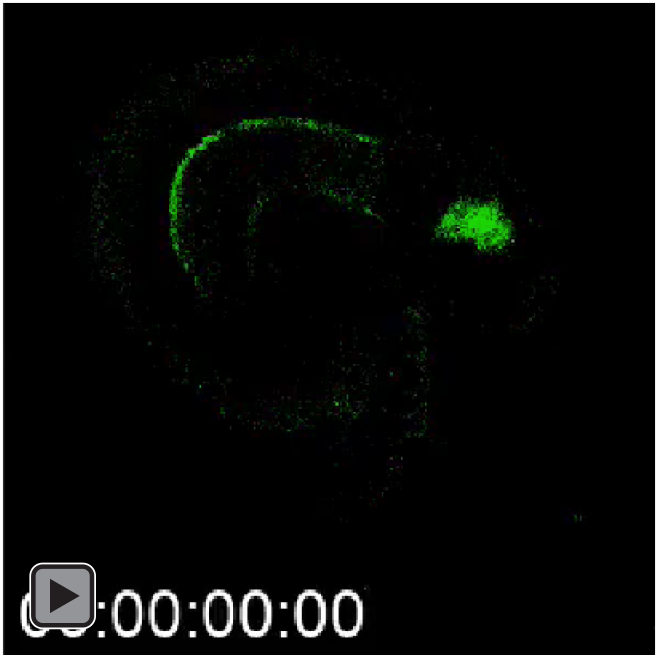
NTA-GFP (green signal) accumulates at the filiform apparatus during pollen tube reception (GFP channel only, same movie as S1).

**Supplementary movie 3.**
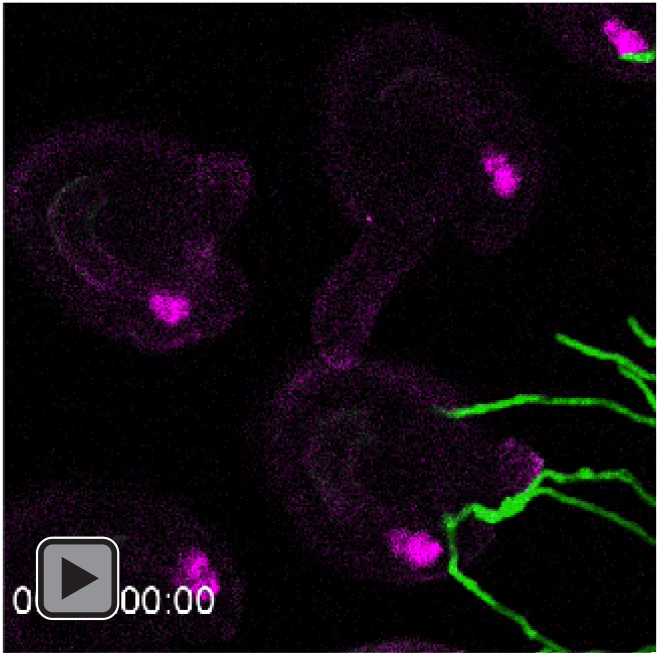
Golgi-mCherry signals (magenta signal) are evenly distributed along the length of the synergid as Lat52::GFP labeled pollen tube (green signal) approaches.

**Supplementary movie 4.**
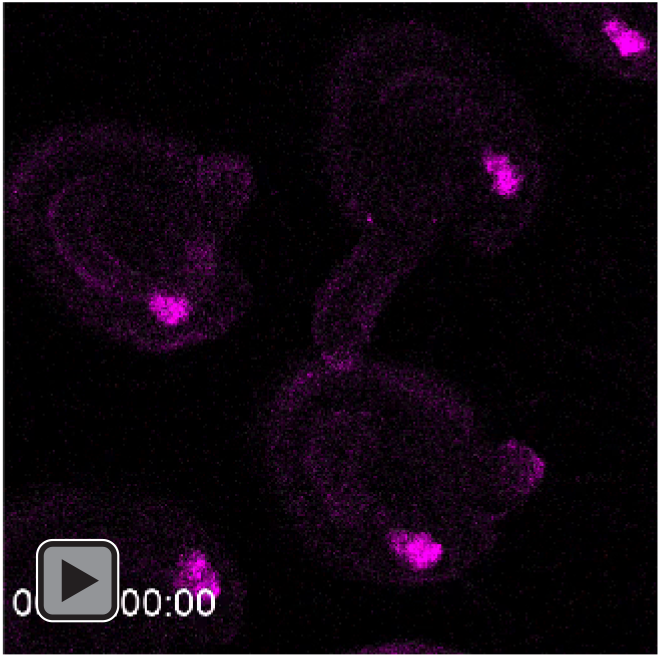
Golgi-mCherry signals (magenta signal) are evenly distributed along the length of the synergid during pollen tube reception (mCherry channel only, same movie as S3).

**Supplementary movie 5.**
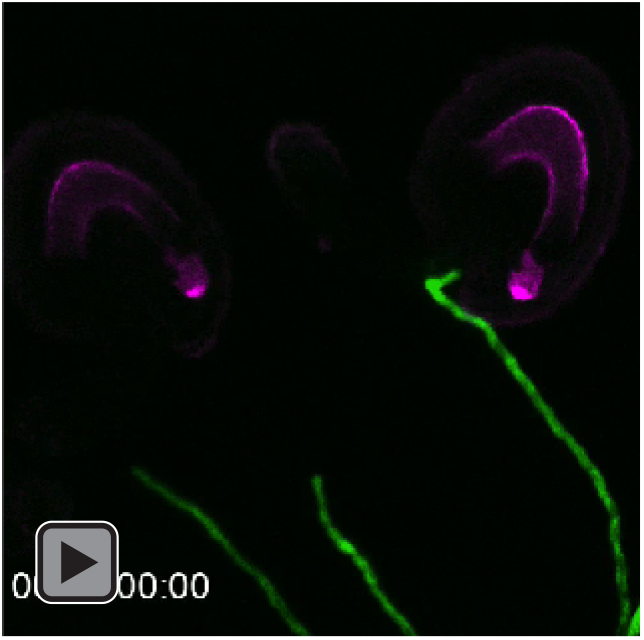
The trans-Golgi marker SYP61-mCherry (magenta signal) is localized toward the micropyle region of synergid cells both before and after pollen tube (green signal) arrival.

**Supplementary movie 6.**
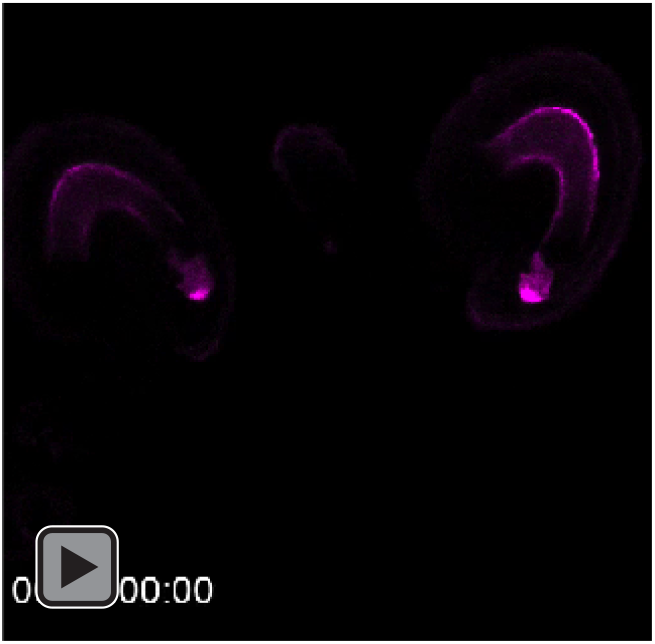
The trans-Golgi marker SYP61-mCherry (magenta signal) is localized toward the micropyle region of synergid cells during pollen tube reception (mCherry channel only, same movie as S5).

**Supplementary movie 7.**
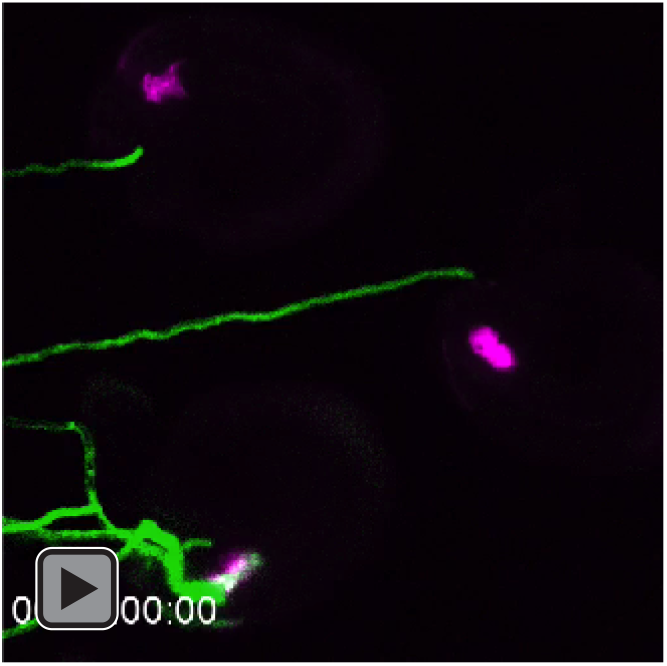
Before and after pollen tube (green signal) arrival, the ER marker SP-mCherry-HDEL (magenta signal) is distributed throughout synergid cells.

**Supplementary movie 8.**
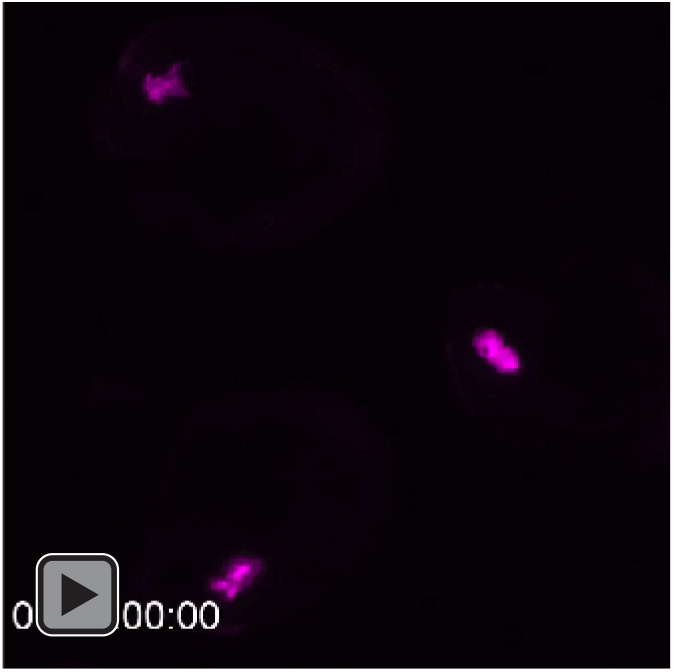
The ER marker SP-mCherry-HDEL (magenta signal) is distributed throughout synergid cells (mCherry channel only, same movie as S7).

**Supplementary movie 9.**
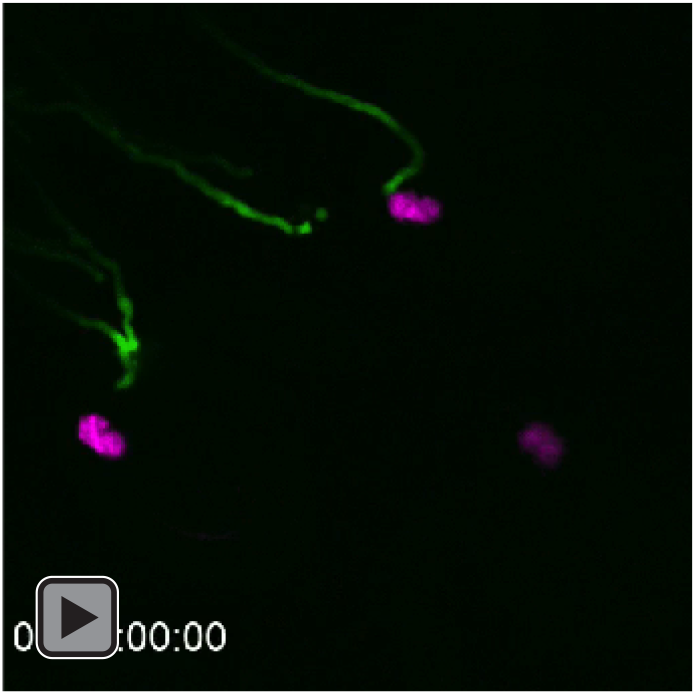
The peroxisome marker mCherry-SLK (magenta signal) does not accumulate at the filiform apparatus after pollen tube (green signal) arrival.

**Supplementary movie 10.**
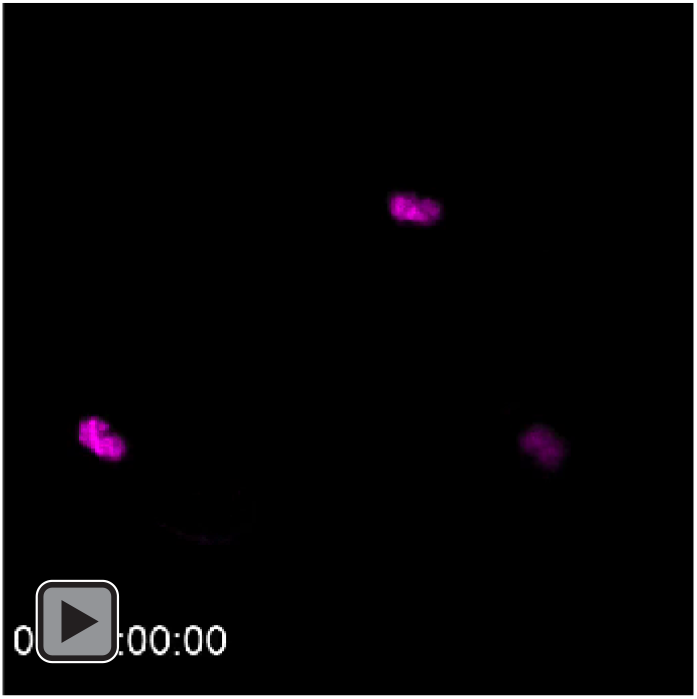
The peroxisome marker mCherry-SLK (magenta signal) does not accumulate at the filiform apparatus during pollen tube reception (mCherry channel only, same movie as S9).

**Supplementary movie 11.**
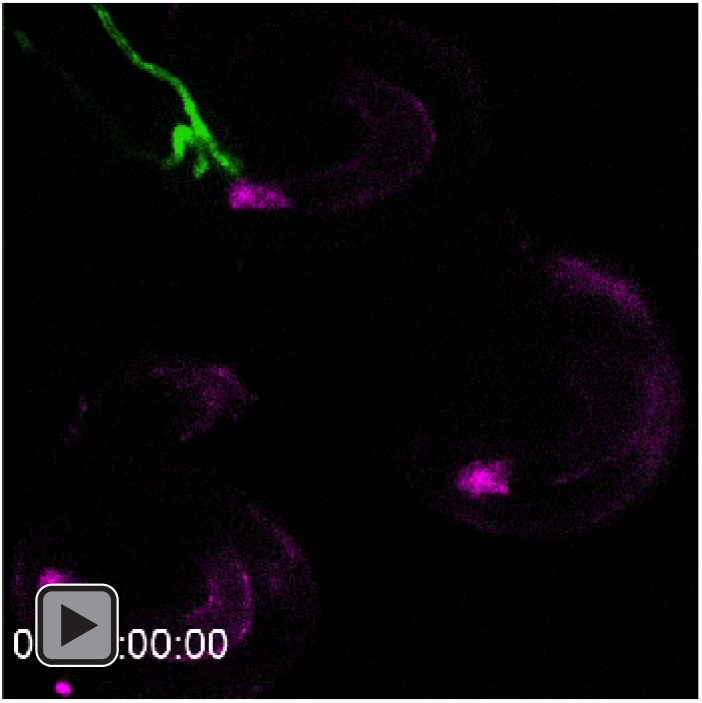
RabA1g-mCherry endosome marker (magenta signal) accumulates at the filiform apparatus in response to pollen tube (green signal) arrival.

**Supplementary movie 12.**
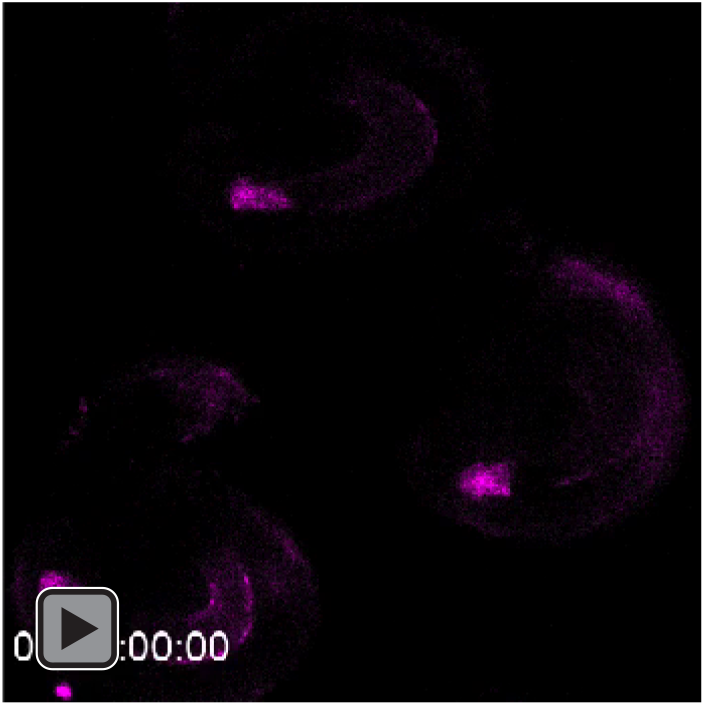
RabA1g-mCherry endosome marker (magenta signal) accumulates at the filiform apparatus in response to pollen tube (green signal) arrival (mCherry channel only, same movie as S11).

## Notes

### Competing Interest Statement

The authors have declared no competing interest.

### Summary of Updates

Manuscript text and figures were edited and new data was added.

